# Loss of virulence of *Botrytis cinerea* mutants defective in phytotoxin production is restored by modifying inoculation medium

**DOI:** 10.1101/2024.10.17.618980

**Authors:** Si Qin, Xiaoqian Shi-Kunne, Jie Chen, Henriek G. Beenen, Yaohua You, Jan A. L. van Kan

## Abstract

Botrydial, botcinic acid and their derivatives are major phytotoxic metabolites produced by the necrotrophic fungal pathogen *Botrytis cinerea*. These phytotoxins are able to induce programmed cell death in the host and thereby promote plant susceptibility to *B. cinerea.* We observed that a *Δbot2Δboa6* double mutant strain, which synthesizes neither botrydial nor botcinic acid, was almost avirulent on tomato leaves when the disease assay was performed using synthetic minimal Gamborg B5 medium. However, virulence of this mutant was restored when the inoculation medium was supplemented with yeast extract. Further virulence assays which compared the double mutant with other multiple mutants using both inoculation media, revealed a prominent contribution of botrydial and botcinic acid to the full virulence of *B. cinerea*. Therefore, we performed an RNA-sequencing experiment to identify *B. cinerea* genes that contribute to the phenotypic switch from an “incompatible” to a “compatible” interaction between tomato and the *Δbot2Δboa6* double mutant. Four genes encoding cell death-inducing effector proteins were upregulated in *B. cinerea* by the addition of yeast extract, and their transcript profiles grouped within a co-expression module that was positively correlated with the compatible interaction. Functional analyses of these effector genes were performed by overexpressing them individually in the *Δbot2Δboa6* background, followed by disease assays with the Gamborg B5 medium without yeast extract.

**Importance:** The grey mould fungus *Botrytis cinerea* is a model for necrotrophic plant pathogens due to its wide host range, economic impact, well-assembled genome, and versatile mechanisms for inducing host cell death during colonization. Botrydial and botcinic acid have previously been characterized as major phytotoxins produced by *B. cinerea*. However, studies from different groups reported variable results regarding the contributions of these phytotoxins to fungal virulence. Here we demonstrate that botrydial and botcinic acid make a prominent contribution to the full virulence of *B. cinerea*, by performing infection assays with mutants that are defective in phytotoxin production and/or multiple cell death-inducing proteins using different inoculation media. This work highlights the pivotal roles of these phytotoxins as compared to other virulence factors, as well as the significant impact of inoculation conditions on compatible and incompatible interactions between the fungus and its hosts.

## Introduction

The grey mould *Botrytis cinerea* is a plant pathogen that can infect more than 1000 host species (1). As a necrotroph, *B. cinerea* kills host cells and feeds on dead tissues after penetrating the plant surface. To induce host cell death, *B. cinerea* secretes a cocktail of cell death-inducing molecules (CDIMs), including phytotoxic secondary metabolites (SMs) and cell death-inducing proteins (CDIPs) (2, 3). The most abundantly produced phytotoxic SMs produced by *B. cinerea* are the sesquiterpene botrydial (BOT) and polyketide botcinic acid (BOA) (4–6). CDIPs secreted by *B. cinerea* and their modes of action are more extensively studied than phytotoxic SMs. Thus far 19 *B. cinerea* secreted proteins were demonstrated to be capable of inducing cell death in at least one plant species (7–11). However, the number of proteinaceous effectors of *B. cinerea* was predicted to be more than 180 (12), and this fungus may secrete more protein effectors functioning in cell death induction than the currently validated number. Phytotoxic SMs and CDIPs contribute collectively to fungal virulence, with a high functional complementarity (7), but the quantitative contribution of each individual CDIM remains to be clarified.

A study by Dalmais et al. (2011) (13) analyzed the roles in virulence of BOT and BOA, either separate or in combination, by testing knockout mutants in key biosynthetic genes. Single mutants defective in the production of either BOT or BOA displayed similar virulence as the wild type (WT) while double mutants that produced neither BOT nor BOA formed ∼50% smaller lesions on French bean (13). However, a recent study described no difference in virulence on tomato leaves between a *Δbot2Δboa6* double mutant generated by CRISPR/Cas-mediated transformation and WT (14). The difference between studies suggests that the role of BOT and BOA in virulence of *B. cinerea* requires further investigation. Commonly, the contribution of (single or multiple) genes to fungal virulence is tested by disease assays in which lesion sizes of a WT recipient fungus are compared with mutants lacking the gene(s) encoding putative virulence factors. Such assays can be executed under different conditions, with variations by different labs either in the inoculation medium, fungal tissue (mycelial plugs or conidia), plant growth conditions or the incubation after inoculation. Although these previous studies both performed assays using conidia suspensions to inoculate tomato leaves, there were differences in the inoculation media, spore density, volume of droplets and in growth conditions for plants (13, 14). We aimed to obtain better understanding of the molecular basis for the fact that different inoculation conditions with the same set of mutants yields such different results. Here, we describe that inoculation of the *Δbot2Δboa6* double mutant in a synthetic medium can result in restriction of the fungus to necrotic spots at the inoculation site (“incompatible interaction”), whereas supplementation of the medium with yeast extract restored the development of expanding lesions (“compatible interaction”). To obtain insights into the mechanisms that regulate such unexpected (binary) outcome of an inoculation, we performed an RNA-seq study. We inoculated the *Δbot2Δboa6* double mutant on tomato leaves in two distinct inoculation media resulting in either compatible or incompatible interactions. As a control, the WT was inoculated in the same media, which in both cases resulted in a compatible interaction on tomato leaves. Inoculated leaves were sampled during the early interaction phases for RNA extraction and sequencing. Resulting data were analyzed to investigate which differences in transcript profiles may underlie the distinction between compatible and incompatible interactions.

## Results

### The importance of BOT and BOA for fungal virulence depends on the inoculation medium

The BOT gene cluster in *B. cinerea* consists of seven genes, including the gene *Bcbot2* which encodes a sesquiterpene cyclase that converts the precursor farnesyl diphosphate (FPP) to presilphiperfolan-8β-ol (15, 16). The BOA gene cluster contains 13 genes (17, 18), of which *Bcboa6* and *Bcboa9* encode polyketide synthases that are key enzymes for BOA biosynthesis (13). We performed infection assays by inoculating tomato leaves with the *Δbot2#1* mutant that cannot produce BOT, the *Δboa6#1* mutant that produces no BOA, and the *Δbot2Δboa6#6* double mutant that produces neither BOT nor BOA. The infection assay was performed using a similar medium to previous studies describing *B. cinerea* infection assays (13, 14). More than 80 % of the spots inoculated with WT B05.10 produced expanding lesions (Fig. 1A, C). The *Δboa6#1* single knockout mutant displayed similar disease incidence and expanding lesion sizes as WT (Fig. 1A, C), suggesting that BOA alone does not make a detectable contribution to virulence on tomato. By contrast, ∼30% of the primary lesions formed by the *Δbot2#1* mutant expanded beyond the inoculation spot, while ∼97% of primary lesions formed by WT did. The expanding lesions that the *Δbot2#1* mutant developed were ∼25% smaller in size as compared to WT (Fig. 1A, C). This observation suggested that production of BOT contributes strongly to the ability to expand beyond the inoculation spot, and to some extent to the expansion rate of lesions. Expanding lesions were barely observed on the leaf half inoculated with the *Δbot2Δboa6#6* double mutant (average disease incidence = ∼6%) (Fig. 1A, C), and the few lesions that expanded were similar in size to the WT, due to the small sample number which hampered proper statistical analysis. These observations indicate that production of BOT or BOA is pivotal for the capacity of *B. cinerea* to cause expanding lesions under these inoculation conditions, with BOT playing a more important role than BOA in virulence.

**Fig. 1.**
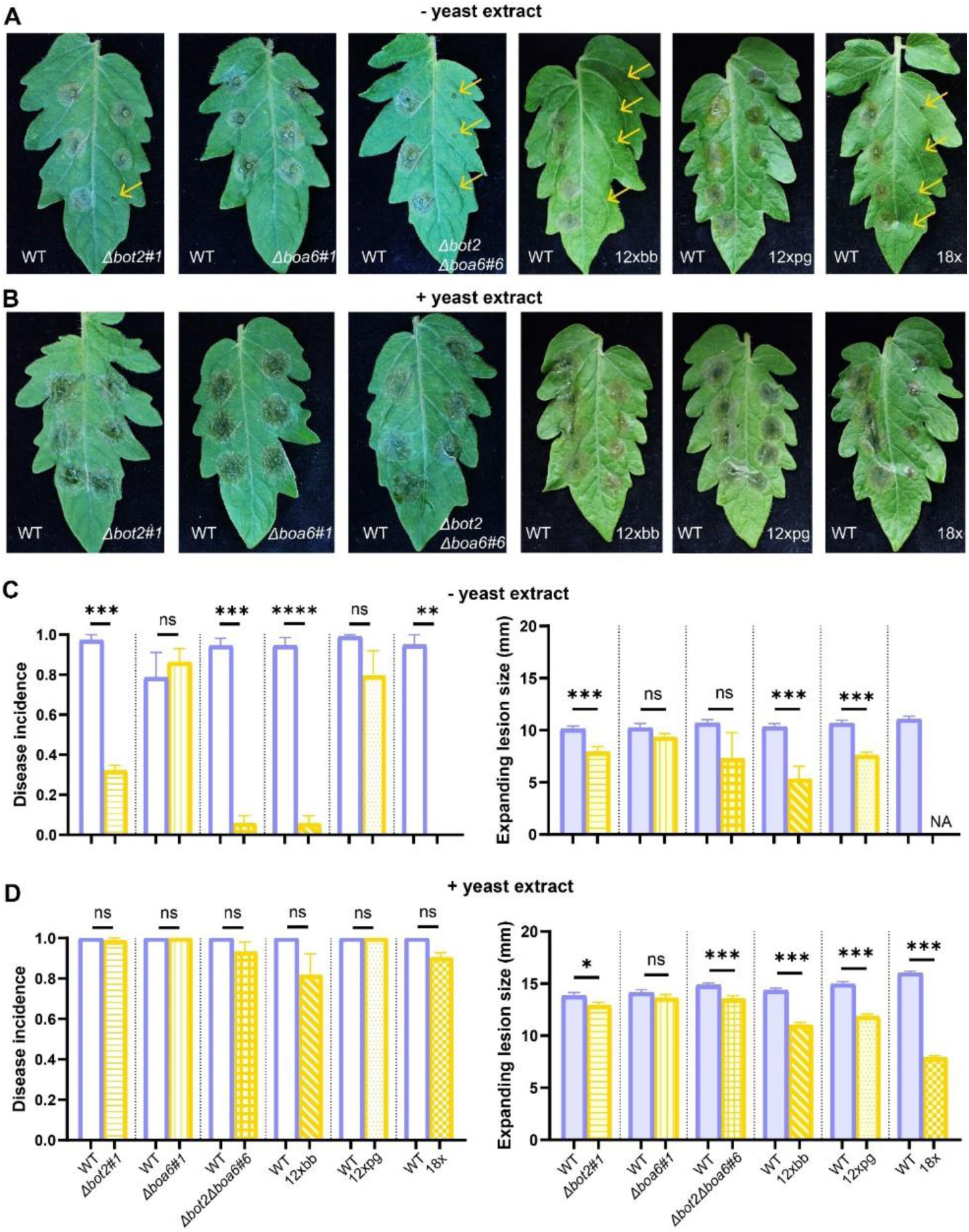
Infection assays to compare the virulence of *Δbot2#1, Δboa6#1* and *Δbot2Δboa6#6* and 12xbb, 12xpg and 18x mutants with WT B05.10 on tomato leaves using two inoculation media. The assays were performed using Gamborg B5 medium without yeast extract (A, C) or with 0.1% yeast extract (B, D). Symptoms of tomato leaflets infected by each mutant strain (on the right of the central vein) compared with B05.10 (on the left of the central vein) were photographed at 3 days post inoculation (dpi) (A, B). Yellow arrows in (A) indicate non-expanding lesions. Bar charts of disease incidences (left) and lesion sizes measured by a digital caliper (right) at 3 dpi, present the means with standard errors from 72 (for *Δbot2#1, Δboa6#1* and *Δbot2Δboa6#6* compared to WT fungus) or 96 inoculations (for 12xbb, 12xpg and 18x mutants compared to WT fungus) collected from three independent experiments (C, D). The virulence assays for *Δbot2#1, Δboa6#1* and *Δbot2Δboa6#6* were performed separately from the experiments for 12xbb, 12xpg and 18x mutants. Disease incidence was calculated as the ratio of the number of expanding lesions to the total number of inoculated spots. Lesions showing a diameter no larger than 2 mm were considered as non-expanding lesions. Statistical analyses were performed by t-test, and the results are shown by either asterisks indicating the significant differences (*p<0.05, **p<0.01, ***p<0.001, ****p<0.0001) or ns, indicating no significance.

When the infection assay was performed using the same fungal strains but with addition of 0.1% yeast extract in the inoculation medium, the capacity of *Δbot2#1* and *Δbot2Δboa6#6* mutants to cause expanding lesions was largely restored and comparable to the WT. Under this modified condition (+y), disease incidence of WT increased to 100% and the lesion size increased by approx. 5 mm compared to the former condition (-y) (Fig. 1B, D). The *Δboa6#1* single knockout mutant displayed similar disease incidence and lesion sizes as WT (Fig. 1B, D). Interestingly, almost all sites inoculated with *Δbot2#1* and *Δbot2Δboa6#6* mutants showed expanding lesions. Lesion sizes of these mutants were slightly, but significantly smaller than those of the WT (Fig. 1B, D). Similarly, 100% disease incidence was observed for WT, as well as the single and double phytotoxin biosynthetic mutants, when inoculation was performed using potato dextrose broth (Fig. S1), a commonly used medium for fungal infection assays.

### BOT and BOA have a larger impact on the virulence of *B. cinerea* than CDIPs

Two recent studies reported that the contribution of CDIMs of *B. cinerea* displays a high level of functional complementarity, as demonstrated by the phenotypes of 12xbb, 12xpg and 18x mutants (in which 12, 12 and 18 *B. cinerea* genes are knocked out, respectively; Table 2) (7, 19). We performed infection assays to compare the virulence of the 12xbb, 12xpg and 18x mutants to WT under inoculation conditions described above. The *Δbot2Δboa6#6* strain was assessed along with these mutants in the same experiments. When no yeast extract was added into the inoculation medium, even fewer spots inoculated with the 12xbb mutant showed expanding lesions as compared to *Δbot2Δboa6#6,* while the 18x mutant was unable to cause even a single expanding lesion on the tomato leaves (Fig. 1). The 12xpg mutant caused expanding lesions at most inoculation spots, although the lesion sizes were significantly smaller than for the WT (Fig. 1). These observations collectively suggested that the deletion of BOT and BOA caused a major reduction in virulence, while the 12 genes missing in the 12xpg played a less pronounced role, as compared to BOT and BOA when tested under this condition (-y). After adding 0.1% yeast extract into the inoculation medium, disease incidences for *Δbot2Δboa6#6*, 12xbb and 18x mutants increased to above 80% and were no longer significantly different from WT (Fig. 1), however there were notable differences in the sizes of expanding lesions. With this inoculation medium (+y), the reduction in lesion sizes of the 12xpg mutant was similar to the *Δbot2Δboa6#6* double mutant, but less pronounced than the reduction in lesion sizes of the 12xbb and 18x mutants (Fig. 1). This observation suggests that the contribution of the phytotoxins BOT and BOA to virulence was quantitatively comparable to the contribution of the 12 CDIP-encoding genes which were knocked-out in the 12xpg mutant. The 18x mutant showed the most pronounced reduction in lesion size among the tested knockout strains (Fig. 1), indicating additive effects of the deletion of *Bcssp2, Bccfem1, Bccdi1* and *Bccrh1* (knocked out in the 18x mutant but not in the other mutant strains) on the virulence of *B. cinerea*. Moreover, based on the reduction in lesion sizes, the ten extra genes deleted in 12xbb, as compared with *Δbot2Δboa6#6*, seemed to contribute to the virulence to a similar extent as the six extra genes deleted in 18x, as compared with the 12xbb mutant (Fig. 1).

### RNA sequencing of compatible and incompatible interactions

It was remarkable that the incompatible interaction between the *Δbot2Δboa6#6* mutant and tomato leaves could be converted into a compatible interaction by supplementation of yeast extract to the inoculation medium. We hypothesized that the modification of inoculation medium altered the transcriptome of the *Δbot2Δboa6#6* mutant, either by suppressing the expression of genes that result in restricting the fungus in the primary necrotic lesion, or by promoting expression of genes that overrule plant resistance mechanisms leading to fungal restriction. We performed an RNA-seq experiment to study differences in the transcriptome of the *Δbot2Δboa6#6* mutant between the incompatible and compatible interactions. Conidia of the *Δbot2Δboa6#6* mutant and WT B05.10 were suspended in either of the Gamborg B5 media without or with yeast extract (indicated by “-y” or “+y”, respectively) and inoculated on tomato leaves, which were sampled at 0, 12, 16 and 24 hpi. These timepoints were chosen as they represent crucial moments in the infection: at 12 hpi, *B. cinerea* has penetrated the leaf surface and colonizes the internal tissue, but host cell death has not yet been initiated; 16 hpi marks the onset of the appearance of necrotic spots on the upper and lower sides of the leaf; 24 hpi marks the initiation of lesion expansion beyond the boundaries of inoculation droplets. Mock-inoculated tomato leaves and *in vitro* cultures (both -y and +y media) of *Δbot2Δboa6#6* mutant and WT at the same time points were included as controls. RNA extracted from these samples was used for sequencing (Table S1). A total of 13,754 *B. cinerea* transcripts were analyzed. Principal Component Analyses (PCA) showed all samples clustered as expected (Fig. S2). Specifically, the biological replicates clustered within time points and all *in vitro* samples and all *in planta* samples clustered together, respectively. Since we were mainly interested in providing an explanation for the yeast extract-induced transition from an incompatible to a compatible interaction with the host plant, we focused our initial analysis on the *in planta* conditions. To examine the influence of adding yeast extract on the fungal transcriptome, we compared the samples that were grown in media -y and +y across all time points. The highest number of DEGs was observed at 24 hpi, reflecting that a large number of genes were affected by yeast extract at the late time points, in both wild type (WT) and mutant samples (Table 1).

**Table 1.** Number of Differentially Expressed Genes (DEGs) in different comparisons.

| Group | *In planta* |  | *In vitro* |  |
| --- | --- | --- | --- | --- |
|  | Up-regulated | Down-regulated | Up-regulated | Down-regulated |
| WT<sup>1</sup>+v<sup>3</sup> (vs) WT-v<sup>4</sup> 0h<sup>5</sup> | 0 | 0 | 0 | 0 |
| WT +v (vs) WT-v 12h<sup>5</sup> | 85 | 103 | 24 | 7 |
| WT +v (vs) WT-v 16h<sup>5</sup> | 50 | 20 | 7 | 24 |
| WT +v (vs) WT-v 24h<sup>5</sup> | 329 | 455 | 120 | 18 |
| △△<sup>2</sup>+v (vs) △△-v 0h | 3 | 3 | 0 | 0 |
| △△ +v (vs) △△-v 12h | 81 | 186 | 60 | 14 |
| △△ +v (vs) △△-v 16h | 129 | 44 | 70 | 20 |
| △△ +v (vs) △△-v 24h | 771 | 513 | 111 | 77 |

<sup>1</sup> Wild type; <sup>2</sup> *Δbot2Δboa6#6* double mutant; <sup>3</sup> With yeast extract; <sup>4</sup> Without yeast extract; <sup>5</sup> 0, 12, 16, or 24 hpi

| Group | *In planta* |  | *In vitro* |  |
| --- | --- | --- | --- | --- |
|  | Up-regulated | Down-regulated | Up-regulated | Down-regulated |
| WT<sup>1</sup>+y<sup>3</sup> (vs) WT-y<sup>4</sup>_0h<sup>5</sup> | 0 | 0 | 0 | 0 |
| WT +y (vs) WT-y_12h<sup>5</sup> | 85 | 103 | 24 | 7 |
| WT +y (vs) WT-y_16h<sup>5</sup> | 50 | 20 | 7 | 24 |
| WT +y (vs) WT-y_24h<sup>5</sup> | 329 | 455 | 120 | 18 |
| △△<sup>2</sup>+y (vs) △△-y_0h | 3 | 3 | 0 | 0 |
| △△ +y (vs) △△-y_12h | 81 | 186 | 60 | 14 |
| △△ +y (vs) △△-y_16h | 129 | 44 | 70 | 20 |
| △△ +y (vs) △△-y_24h | 771 | 513 | 111 | 77 |

<sup>1</sup> Wild type; <sup>2</sup> *△bot2△boa6#6* double knockout mutant; <sup>3</sup> With yeast extract; <sup>4</sup> Without yeast extract ;<sup>5</sup> 0, 12, 16, or 24 hpi

**Table 2.**
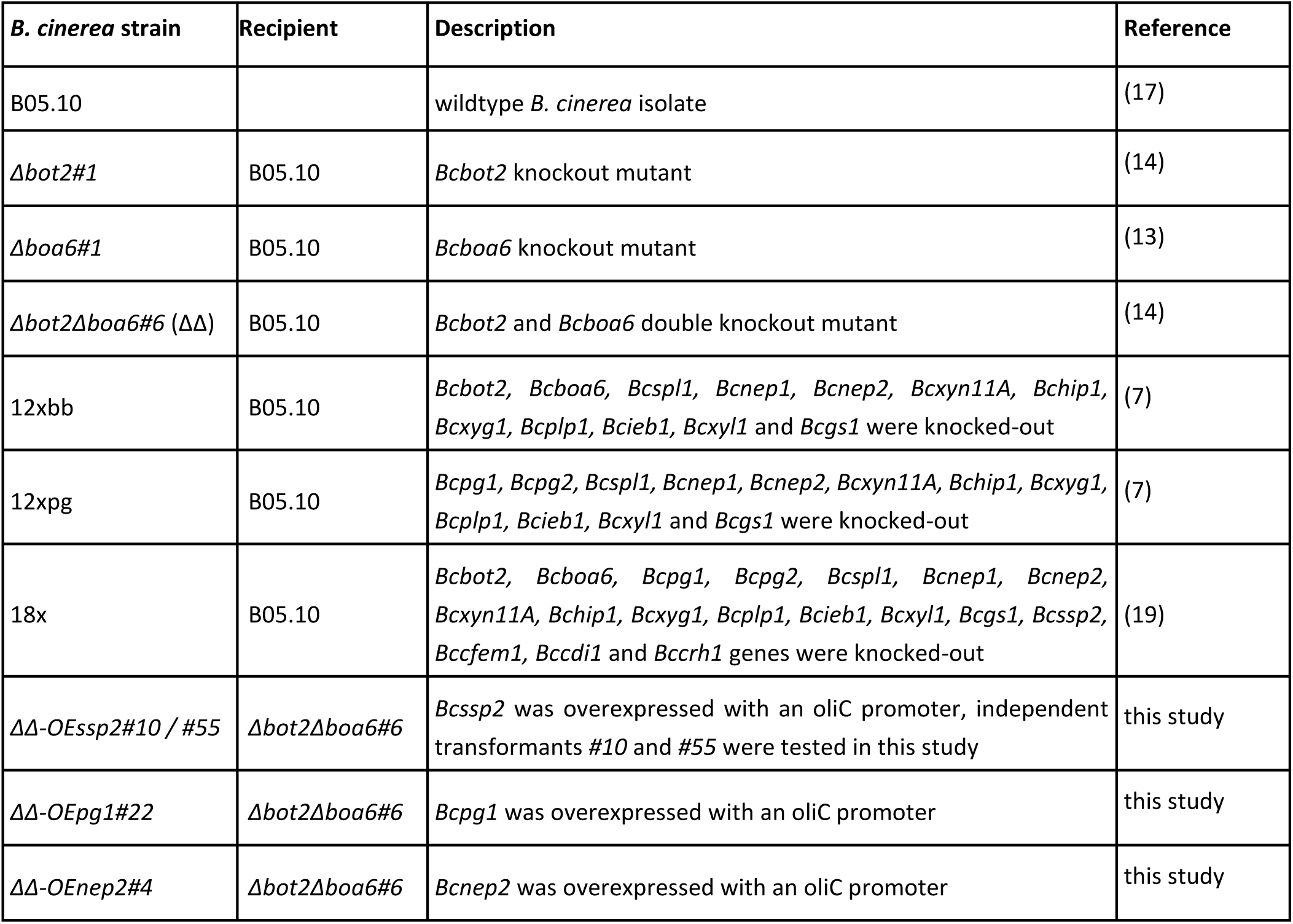
*B. cinerea* strains used in this study.

A large number of the DEGs at 24 hpi (1491) overlaps between WT and mutant (Fig. 2), which indicates that the fungal strains were similarly affected by the yeast extract. The up-regulated genes are mostly enriched in GO terms (p < 0.05) related to gene transcription and protein synthesis (Fig. S3). Enriched GO terms detected in the set of down-regulated genes are involved in iron/copper homeostasis. Moreover, *Bcatg13, Bcatg2* (20) and *Bcatg9* that are involved in autophagy (ATG) were also down-regulated. ATG includes a set of programmed cell developmental changes that occur during cellular remodeling and serves as an adaptive response during nutrient starvation (21). In stress conditions ATG may activate a programmed cell death cascade. The upregulation of transcripts of ATG genes in the mutant upon inoculation without yeast extract at 24 hpi coincides with the restriction of lesion outgrowth. There are also 722 and 289 DEGs that were exclusively differentially expressed in the mutant or wild type fungus at 24 hpi, respectively. Enriched GO terms were found only in uniquely downregulated wild type and uniquely upregulated mutant gene lists. The enriched GO terms of the genes that were uniquely up-regulated in the mutant, are related to rRNA processing, protein translation, gene expression, oxidation-reduction, transmembrane transport (Fig. S4).

**Fig. 2.**
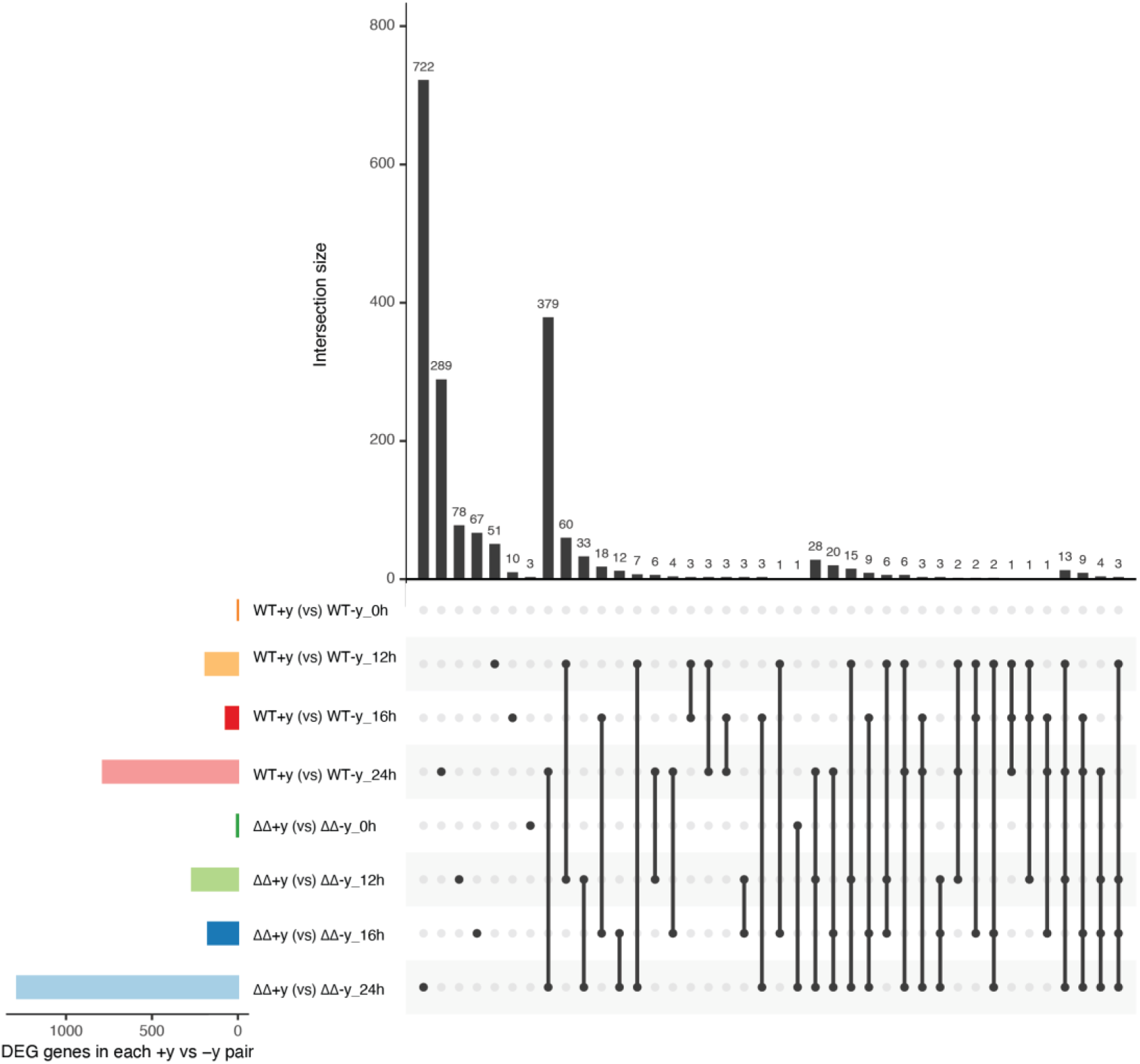
UpSet plot summarizing differentially expressed genes (DEGs) of either wild type or double mutant between samples with and without yeast, at different time points. The vertical bar plot reports the intersection size, the dot plot reports the set participation in the intersection. Links between different samples show overlapped DEGs between or among compared samples. The bottom left horizontal bar graph shows the total number of DEGs per compared group (set size).

### Co-expression network analysis reveals gene clusters correlated with compatible interaction

In order to determine which genes showed similar expression patterns and are co-regulated across all conditions, a co-expression network was created using WGCNA. A total of 25 modules of co-expressed genes was obtained, of which the “Grey” module was the residual module, containing all genes that did not show significant correlation in expression profile with genes in other modules (Fig. S5). Evaluation of interconnections between all 25 modules revealed that each module correlated with its own the best, as to be expected (Fig. S6).

To analyze which modules might be functionally involved either in the response to the presence of yeast extract in the inoculation medium, or in the outcome of the interaction between *B. cinerea* and tomato, we correlated genes of each module with their experimental variables. There was no significant correlation between any module and the presence of yeast extract (Fig. 3). By contrast, nine modules showed significant (positive or negative) correlations with the outcome of the infection, i.e. either with a compatible interaction or an incompatible interaction, respectively.

**Fig. 3.**
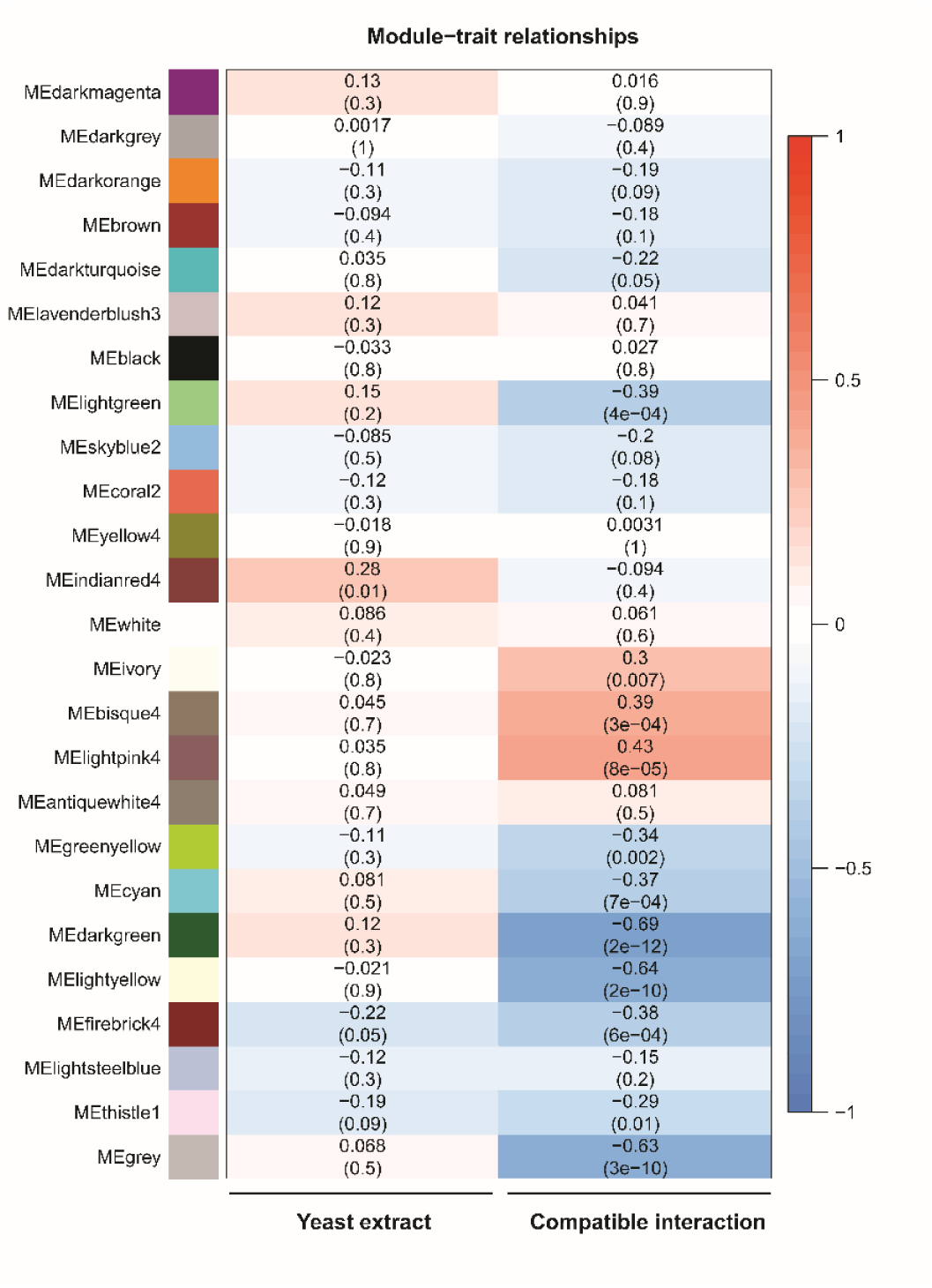
Heatmap showing the correlations of co-expression modules with the presence of yeast extract in the inoculation medium, or in the outcome of the interaction between *B. cinerea* and tomato. On the Y-axis are the co-expression modules, with both the names and colors assigned by the WGCNA algorithm. In the heatmap, Pearson correlations with the experimental variables (the presence of yeast extract and outcome of the interaction) are shown, respectively, left and right. Dark red colors indicate a high positive correlation, and dark blue colors indicate a high negative correlation with a condition. Lighter values indicate lower (positive or negative) correlation. For each module, the top line provides the correlation coefficient r, while the lower line (in a bracket) provides the p-value for significance.

Three modules were positively correlated to the compatible interaction (conditions in which the lesions expanded), and their expression peaks were more pronounced *in planta* than *in vitro* (Fig. 4). The “bisque4” and “light pink4” modules contain genes with transient peaks in transcript levels at 12 hpi and 16 hpi, respectively, whereas genes in the “ivory” cluster showed steady transcript levels at 12 and 16 hpi, followed by a strong increase at 24 hpi, especially in the compatible interactions. Interestingly, all three modules are enriched in proteins with a signal peptide for secretion (p < 0.01). It should be noted that two modules include genes encoding known CDIPs. Module “bisque4” contains the *Bcnep1, Bcxyg1,* and *Bcplp1* genes, while “ivory” contains the genes *Bcnep2, Bcxyn11A, Bcpg1* and *Bcssp2* (Data S1). Surprisingly, the module “lightyellow” that showed negative correlation to the compatible interaction contains even more of such CDIP-encoding genes, specifically *Bcspl1, Bchip1, Bcxyl1, Bcgs1, Bcbot2* and *Bccrh1*.

**Fig. 4.**
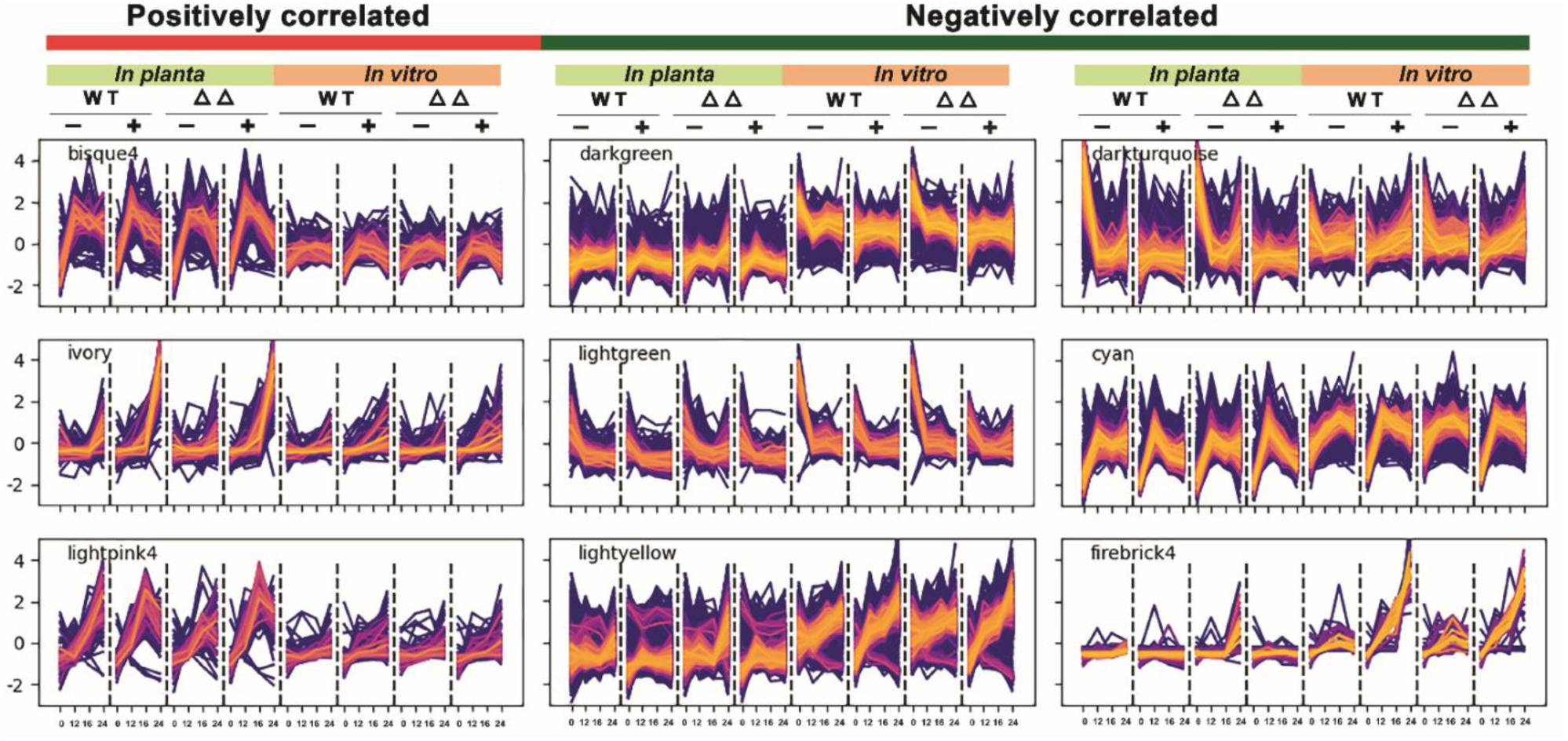
Expression profiles of co-expression modules that show significant (positive or negative) correlations to the compatible interaction between *B. cinerea* and tomato. Each sub-plot contains the full profile of all genes in a module, across all eight conditions. From left to right on the X-axis contains *in planta* and *in vitro* conditions (with the time points 0, 12, 16 and 24 hpi): wild type without yeast extract, wild type with yeast extract, double mutant without yeast extract, double mutant with yeast extract. On the Y-axis, the standardized gene expression (z-score) is shown. Each profile is colored according to the assigned module membership value, defined as the correlation with the first principal component of the module profiles (module Eigengene). Lighter colors indicate greater module membership values.

We examined in detail the three modules positively correlated with the compatible interaction, focusing on their expression profiles, gene content and GO enrichment. Despite each showing a positive correlation to the compatible interaction, these three modules were quite distinct in their enriched GO terms (Fig. S7). The “bisque4” module contains 245 genes that showed a transient peak in expression *in planta* at 12 hpi, followed by a slight decline, while *in vitro* transcript levels remained fairly constant (Fig. 4). “Bisque4” is enriched in genes involved in mitochondrial activity and splicing. The “lightpink4” module contains 85 genes that showed a transient peak in expression *in planta* at 16 hpi for the +y samples, while transcript levels increased in the -y samples (Fig. 4). The “Lightpink4” has no enriched GO terms, but the gene list contains CAZyme-encoding genes, including pectin methylesterase genes, as well as cytochrome P450-encoding genes, including several from the BOA gene cluster. Genes in the “ivory” cluster (118 genes) showed a continuous increase in transcript levels over infection time points, with the increase more pronounced in the +y samples as compared to -y samples. GO terms enriched in the “ivory” cluster are associated with peptidase and hydrolase activities.

We also examined modules that were correlated with the incompatible interaction, displaying expression profiles associated with a failure to cause expanding lesions (Fig. 3). The “darkgreen” module (1604 transcripts) is typified by generally stable *in planta* transcript levels that increase at 24 hpi exclusively in the incompatible interaction. The genes in this module are enriched for GO terms related to gene expression, chromatin organization, splicing, mitotic cell cycle, cell division and DNA repair. The “darkturquoise” module (1811 transcripts) is characterized by high *in planta* transcript level in the absence of yeast extract, particularly at 0 hpi and 24 hpi, with a steep decline at 12 hpi and 16 hpi, while the expression in the presence of yeast extract remains more stable at all time points. This module contains six genes encoding light-dependent transcription factors (TFs), several light sensors, proteins from the Velvet complex, both phospholipase C proteins, as well as five polyketide synthases. The “lightyellow” module (2504 transcripts) is marked by increased *in planta* transcript levels in the absence of yeast extract at 24 hpi, which are higher in the mutant than in the wild type. The only enriched GO terms for this module are related to protein translation. The “lightgreen” module (372 transcripts) is typified by a decline of *in planta* transcript levels over time in the three compatible interactions, while the expression increases at 24 hpi in the incompatible interaction. This module contains RNA polymerase I transcription initiation factors, two HHK histidine kinases and other signaling proteins, bicarbonate transporters and other H+ antiporters. The “cyan” module (867 transcripts) is characterized by a peak in expression at 12 hpi, which drops at 16 hpi. Only in the inoculation medium with yeast extract, transcript levels continue to decrease at 24 hpi. This module contains proteins of the proteasome complex, ATPase complex, as well as proteins involved in cytoskeleton binding, glycosylation and Golgi vesicle transport. The “firebrick4” module (37 transcripts) is typified by stable *in planta* transcript levels that increase at 24 hpi exclusively in the *Δbot2Δboa6#6* mutant without yeast extract, leading to an incompatible interaction. The module contains five copper transporter genes, a copper-binding protein and a copper metallochaperone, suggesting that the fungus experiences a depletion of copper during the incompatible interaction.

### Overexpressing single CDIPs cannot restore the pathogenicity of *Δbot2Δboa6*

Based on phenotypic observations of the mutants in Fig. 1, and the presence of CDIPs in co-expression modules positively correlated with a compatible interaction, we closely examined the expression profiles of CDIP-encoding genes that are deleted in the 18x mutant. A heatmap was generated to compare transcript levels in WT B05.10 and the *Δbot2Δboa6#6* mutant, both *in vitro* and during leaf infection, in the absence or presence of yeast extract (Fig. 5).

**Fig. 5.**
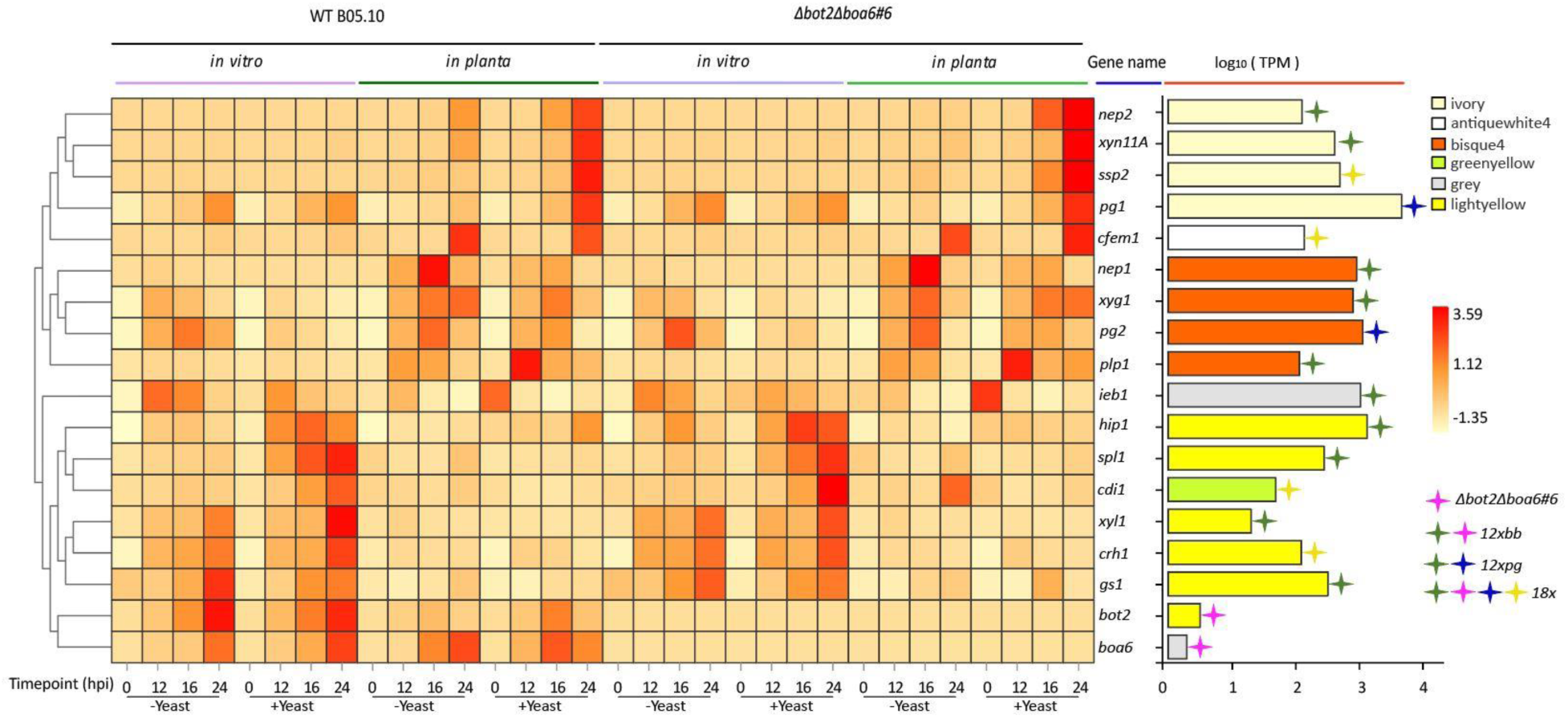
Transcript levels of 18 genes in the WT B05.10 and *Δbot2Δboa6#6*, represented by a heatmap generated by Z-score calculation (left) and the mean log_10_ (TPM) of the gene across all timepoints (right) both during *in vitro* growth and during infection. The genes knocked-out in *Δbot2Δboa6#6*, 12xbb, 12xpg and 18x mutants were marked by stars with corresponding colors, for which the legend is at the right bottom corner of the figure. The bar in the mean log_10_ (TPM) panel is filled with a color representing the module in which the gene grouped according to the WGCNA analysis.

Expression patterns of all 16 genes except for *Bcbot2* and *Bcboa6* were similar between WT B05.10 and *Δbot2Δboa6#6.* Notably, *Bcssp2*, *Bcnep2*, *Bcxyn11A* and *Bcpg1*, members of the “ivory” module, shared a similar transcript profile characterized by their upregulation during tomato infection upon the addition of yeast extract to the inoculum. *Bcssp2* was of particular interest, as it is one of four genes deleted in the 18x mutant but not in the 12xbb or 12xpg mutants, while *Bcnep2*, *Bcxyn11A* and *Bcpg1* were deleted in the 12xpg mutant (Fig. 5). Since the 12xpg mutant was more virulent than *Δbot2Δboa6#6* in the absence of yeast extract and displayed similar virulence as *Δbot2Δboa6#6* in the presence of yeast extract (Fig. 1), we hypothesized that *Bcssp2* plays a more important role in virulence under these conditions than *Bcnep2*, *Bcxyn11A* or *Bcpg1*. Among the genes deleted only in the 18x mutant, *Bcssp2* was the only gene that was upregulated by addition of yeast extract; the transcript level of *Bccfem1* remained similar with or without yeast extract, while *Bccdi1* and *Bccrh1* transcripts were downregulated when yeast extract was added.

To assess whether overexpression of *Bcssp2* in the *Δbot2Δboa6#6* mutant could restore virulence without addition of yeast extract, by compensating for the absence of BOT and BOA production, fungal transformants were generated, using the *Δbot2Δboa6#6* mutant as a recipient, with *Bcssp2* under control of a constitutive promoter. The virulence of *Δbot2Δboa6-OEssp2* (*ΔΔ-OEssp2*) mutants was compared with the recipient using inoculation medium lacking yeast extract. Although the potential contribution of *Bcnep2*, *Bcxyn11A* and *Bcpg1* to the virulence of *B. cinerea* was expected to be less significant than *Bcssp2*, these genes were grouped in the same module based on their expression profile. Therefore, transformants were also generated to separately overexpress either *Bcnep2*, *Bcxyn11A* or *Bcpg1* in the *Δbot2Δboa6#6* recipient (Fig. S8). Unexpectedly, *ΔΔ-OExyn11A* transformants showed growth retardation during *in vitro* growth and were eliminated from infection assays. Two transformants of *ΔΔ-OEssp2*, one transformant of *ΔΔ-OEpg1*, and one for *ΔΔ-OEnep2* were inoculated on tomato leaves in medium lacking yeast extract to compare their virulence with the recipient strain *Δbot2Δboa6#6*. None of the overexpression transformants caused a higher proportion of expanding lesions than *Δbot2Δboa6#6* when inoculated without yeast extract (Fig. 6). Thus, overexpressing *Bcssp2, Bcnep2* or *Bcpg1* individually cannot compensate for the defect in pathogenicity of the *Δbot2Δboa6#6* mutant.

**Fig. 6.**
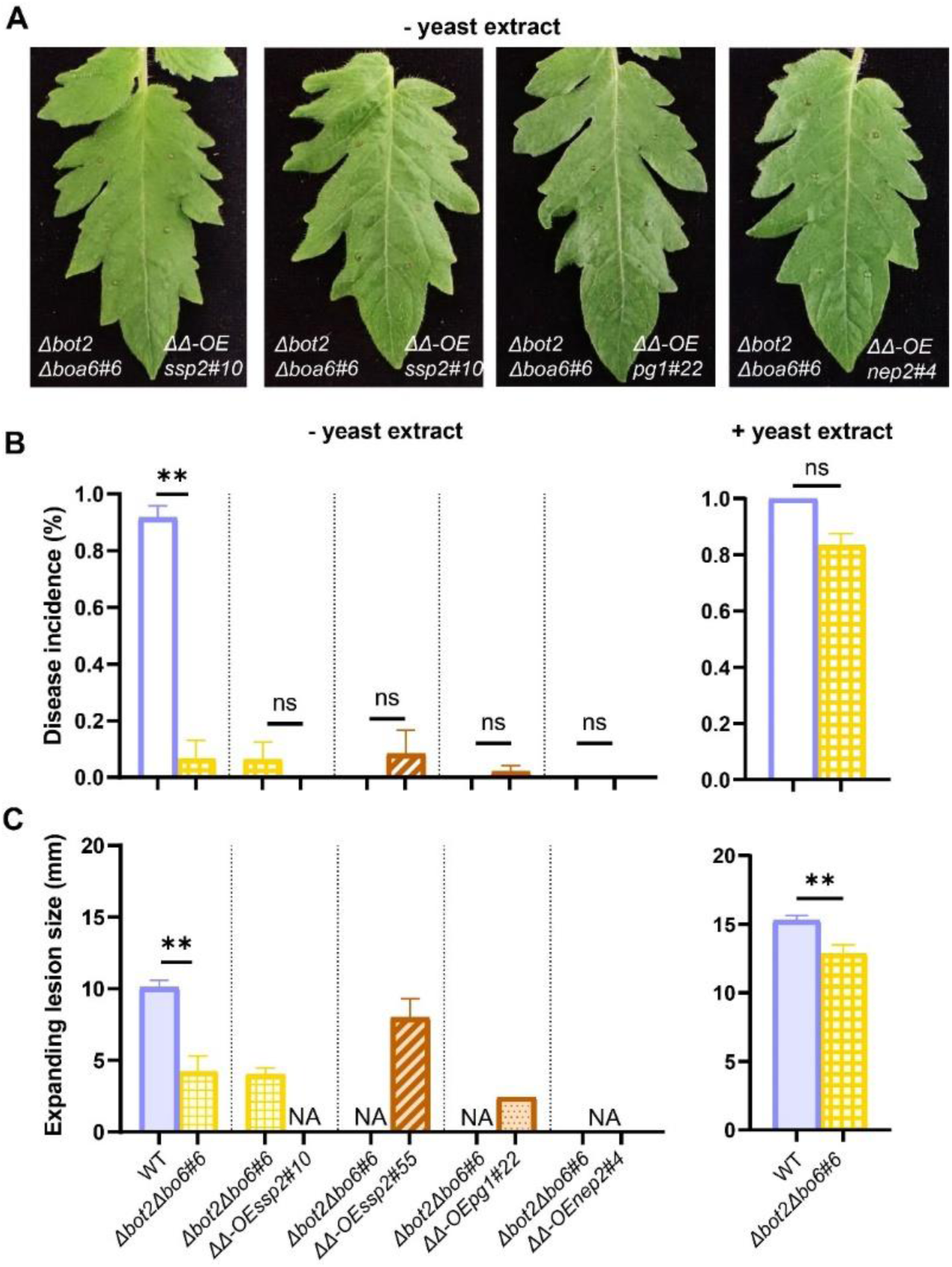
Infection assays to compare the virulence of overexpression mutants to the recipient strain *Δbot2Δboa6#6* on tomato leaves using a Gamborg B5 medium without yeast extract. The comparisons between WT B05.10 and *Δbot2Δboa6#6* were also inoculated using both medium without and with yeast extract in the same experiments, showing similar results as shown in Fig. 1. (A) Symptoms of tomato leaflets infected by each overexpression mutant strain (on the right of the central vein) compared with *Δbot2Δboa6#6* (on the left of the central vein) were photographed at 3 days post inoculation (dpi). (B) Bar charts of disease incidences of inoculations performed using the medium without yeast extract (left) and the disease incidences of WT B05.10 and *Δbot2Δboa6#6* inoculated with the medium with yeast extract in the same experiments (right), which are represented as means with standard errors from 48 datapoints collected from two independent experiments. (C) Bar charts representing expanding lesion sizes upon inoculations using the medium without yeast extract (left) and with yeast extract (right), measured by a digital caliper at 3 dpi. The total number of expanding lesions caused by each of the mutant strains, which were inoculated with the medium without yeast extract, was no more than 5 from these two experiments. The inoculations of mutant strains causing no expanding lesion are indicated as “NA” in the chart on the left. Statistical analyses were performed by t-test, showing no significance (ns) or significant difference (**p<0.01) in disease incidence and expanding lesion size between the mutant and the recipient strain (*Δbot2Δboa6#6* for overexpression mutants or WT B05.10 for the *Δbot2Δboa6#6* mutant).

## Discussion

### The occurrence of an incompatible interaction between a necrotrophic fungus and its host

Most fungal pathologists will by definition consider a gene to encode a virulence factor if a knock-out mutant strain produces lesions with significantly smaller lesion size as compared to the wild type. This definition of a virulence factor is, however, neglecting the contribution of inoculation medium to the eventual disease development. The results of this study illustrate the significant impact of the composition of inoculation medium on disease incidence resulting from inoculation with *B. cinerea* mutants, but not the wild type. The mutants retained their capacity to cause primary necrotic lesions, however, these primary lesions did not expand beyond the inoculation site, resulting in restriction of fungal outgrowth that reflects an “incompatible interaction”. The terms “compatible” and “incompatible” interaction are commonly used for interactions of plants with biotrophic microbial pathogens (22). In that case the outcome of an interaction is often determined by gene-for-gene interactions of pathogen effectors, acting as avirulence factors, with cognate receptors in the host plant that recognize the effector and trigger hypersensitive cell death and resistance (23, 24). We are aware that it appears unusual to use the term (in)compatible for the infection of a necrotrophic fungus, and especially in the context of a fungal mutant in combination with the composition of the inoculation medium. We recently adopted the same terminology to describe the interaction between *B. cinerea* WT strain B05.10 and the wild tomato relative *Solanum habrochaites* in which the sucrose concentration in the inoculum markedly affected the disease incidence (25).

In our experiments with *Δbot2#1* and *Δbot2Δboa6#6* mutants, as well as the 12xbb and 18x mutants, the addition of yeast extract to the inoculum restored disease incidence to the level of the WT B05.10 (“compatible interaction”) although lesion sizes caused by the mutants at 3 dpi were smaller as compared to WT B05.10. The results from virulence assays using both media indicated a more pronounced contribution of BOT and BOA to the virulence of *B. cinerea* than previously reported (13, 14). Leisen et al. (2020) (14) reported that the lesion size caused by the *Δbot2Δboa6#6* mutant was similar to that of WT B05.10, using the same Gamborg B5 medium as in this study (without yeast extract) but using for the infection assay a lower spore density combined with a larger droplet size (the total number of conidia in each droplet was similar between both studies). We also observed that the *Δbot2Δboa6#6* mutant caused expanding lesions on tomato leaves using as the inoculation medium PDB (Fig. S1), which differs in the type and concentration of nutrients from the Gamborg B5 media (-y/+y). We are not aware of any previous study on *B. cinerea* or any other fungus where such seemingly small variation in experimental procedures leads to opposite outcomes of disease development (incompatible versus compatible interaction). Hopefully, these observations will encourage the research community to more thoroughly consider the methodology used for testing the virulence of knockout mutants in *B. cinerea* and possibly other fungi.

These results inspired us to perform an RNA-seq strategy to acquire insights into the “compatible” or “incompatible” interaction of the *Δbot2Δboa6#6* mutant (but not WT B05.10) with tomato Moneymaker leaves and into the effect of adding yeast extract to the inoculum. We considered that restoration of compatibility for the *Δbot2Δboa6#6* mutant by the addition of yeast extract could be mediated by its influences on the transcript levels of fungal genes that govern crucial stages of the infection process, either the primary lesion induction or the subsequent lesion expansion. In order to study these processes with as few experimental variables as possible, we performed an RNA-seq experiment using two media (-/+ yeast extract) as the only difference in the inoculation condition to investigate the molecular mechanisms mediating the transition from an incompatible to a compatible interaction. The focus of our analysis was on the *B. cinerea* gene expression in three (early) phases of the infection: penetration of the host surface (12 hpi), initiation of primary cell death (16 hpi) and onset of lesion expansion (24 hpi).

### The role of yeast extract

So, what could be the physiological impact of yeast extract on a fungus during the infection process on a host plant? Yeast extract is a concentrate of the water-soluble portion of autolyzed *Saccharomyces cerevisiae* cells and provides water-soluble vitamins, amino acids and small peptides to the medium formulation. The Gamborg B5 medium itself consists of minerals, vitamins, and it contains both nitrate and ammonium as nitrogen sources but no carbon source. We supplemented the medium with 25 mM glucose as the carbon source. The Gamborg B5 medium with glucose suffices to allow *B. cinerea* germination and it enables the WT strain B05.10 to achieve primary lesion induction and subsequent expansion of the lesion (“compatible interaction”) in the absence of supplemented yeast extract. However, yeast extract provides amino acids that enable the fungus to accelerate protein biosynthesis, as it sidesteps the requirement to first synthesize an entire set of amino acids from primary nitrogen and carbon sources in the medium. The immediate availability of (a small amount of) amino acids would give *B. cinerea* a head start that can accelerate the production of proteins and metabolites by a few hours, which stimulates host penetration and the subsequent induction of a primary lesion. Although the WT B05.10 did not require yeast extract to achieve a high disease incidence, the addition of yeast extract to the inoculum resulted in a larger lesion size at 3 dpi (Fig. 1), which was presumably facilitated by faster, or more effective, activation of pathogenicity-related mechanisms and thereby enabled the fungus to proceed to the lesion expansion phase at an earlier timepoint. When WT B05.10 was inoculated in Gamborg B5 medium lacking yeast extract, the development of primary lesions was commonly observed from 16 hpi onwards but expansion of primary lesions occurred only beyond 24 hpi. The addition of yeast extract accelerated the onset of expansion, as the lesions were clearly larger than the inoculation droplet at 24 hpi.

### Primary lesion induction and lesion expansion are distinct processes

We propose that a successful infection by *B. cinerea* involves two aspects, which may be governed by distinct fungal virulence factors, and which require that the fungus achieves certain milestones at specific time points. It is not merely sufficient to trigger plant cell death by producing a cocktail of CDIMs, but these must be produced and operate timely to prevent the host plant from mounting defense responses that otherwise restrict fungal outgrowth. In this context it is important to note that the primary lesions that were produced in the incompatible interaction by the *Δbot2Δboa6#6* mutant remained unable to expand once the restriction of lesion outgrowth was observed, regardless how long the incubation was continued. Apparently, the plant response that restricted the expansion provided an effective, absolute host resistance response which was achieved around the 24 hpi benchmark. We thus hypothesize that the lack of BOT and BOA production caused such significant delay in the capacity of the fungus to induce host cell death in a timely manner, that it allowed the plant to mount an effective resistance. This highlights the importance of time in the interaction.

*B. cinerea* expresses different genes at the stages of primary lesion development and lesion expansion. One example is provided by the family of cell death-inducing NLP proteins that contains two members, BcNEP1 and BcNEP2. *Bcnep1* is expressed from 8 hpi onwards and reaches a peak at 12-16 hpi, coinciding with the appearance of primary lesions (26). *Bcnep1* transcript levels decline sharply at 24 hpi coinciding with a strong increase in *Bcnep2* transcript levels that remain high while lesions continue to expand (26). The cell death-inducing capacity of BcNEP1 protein is stronger than of BcNEP2 and is notably faster (4-8 h after protein infiltration) than for BcNEP2 (∼24 hpi) (26, 27). These results support the hypothesis that primary lesion induction and lesion expansion are distinct processes, involving distinct fungal proteins. In addition, You et al. (2024) (28) reported that during infection on tomato, *B. cinerea* expresses the *BcTom*1 gene encoding a ß-xylosidase that hydrolyses the glycoalkaloid α-tomatine and thereby inactivates its antifungal activity. The expression of *BcTom*1 is low in absence of α-tomatine and rapidly induced as soon as the fungus comes into contact with α-tomatine. As α-tomatine is located inside the vacuoles of tomato cells, contact of *B. cinerea* with α-tomatine only occurs when host cells are damaged by cell death induction. The expression of the *BcTom*1 gene can thus serve as a marker for the timing of host cell death in the *B. cinerea*-tomato interaction (29). The expression of *BcTom*1 in this study revealed a different timing between wild type and *Δbot2Δboa6#6* mutant and between inoculation media. In the absence of yeast extract, both wild type and *Δbot2Δboa6#6* mutant at 12 hpi showed similar *BcTom*1 transcript levels that were equal to the 0 hpi level, indicating that the fungus did not yet respond to α-tomatine. When yeast extract was added, the expression already increased at 12 hpi by 4-fold and 7-fold in the *Δbot2Δboa6#6* mutant and wild type, respectively. This observation indicates that host cell death was induced and primary lesion development was likely in progress. *BcTom*1 transcript levels continued to increase in the subsequent time points. The transcript level in the inoculation without yeast extract at 16 hpi was similar to the level of the inoculation with yeast extract at 12 hpi, both for the *Δbot2Δboa6#6* mutant and wild type, suggesting that the addition of yeast extract accelerated the host cell death induction by approximately 4h. At 24 hpi, profiles of the two inoculation conditions diverged notably. *BcTom*1 transcript level for inoculation of the *Δbot2Δboa6#6* mutant without yeast extract was similar between 16 hpi and 24 hpi, while it doubled for the wild type under the same conditions. By contrast, transcript levels for inoculation of the *Δbot2Δboa6#6* mutant and the wild type inoculated with yeast extract were similar at 24 hpi. In all, these results indicate that yeast extract could accelerate the host cell death induction for both *Δbot2Δboa6#6* mutant and wild type, which therefore advanced the timing of the lesion expansion stage. The shorter primary lesion induction stage might have led to successful infection by the *Δbot2Δboa6#6* mutant when yeast extract was supplemented, as the host plant was likely unable to activate effective resistance responses in such short period of time.

### Attempts to restore a compatible interaction in absence of yeast extract by overexpression of CDIPs

Neither of the four CDIP-encoding genes tested (*Bcssp2, Bcnep2, Bcxyn11A* or *Bcpg1*) could, individually, restore the virulence of *Δbot2Δboa6#6* in the absence of yeast extract, which can be explained by three hypotheses. Firstly, additional single (as yet unknown) CDIMs from the ivory module are involved in stimulating primary lesion expansion and thereby overcome the lack of BOT and BOA production. Secondly, multiple previously described CDIPs might jointly restore pathogenicity of the *Δbot2Δboa6#6* mutant via synergistic action. To validate this hypothesis, it may be useful to generate overexpression mutants in which multiple genes are overexpressed, either by using strong constitutive promoters or by identifying a TF that specifically governs the expression of genes in the ivory module and generating a transformant expression a constitutively active form of such a TF. At present, it is unclear which TF(s) regulate the yeast inducible expression of CDIPs, but recent studies on *B. cinerea* transcriptional networks (30) may help to identify TFs that control the expression of virulence factors. Lastly, considering that the successful infection of a host by *B. cinerea* requires milestones to be achieved by the fungus at specific time points, *B. cinerea* should induce host programmed cell death (PCD) at a proper intensity and timing. As proposed in (3), the early phase of the *B. cinerea* – host interaction resembles a “biotrophic infection” (8-16 hpi), during which the fungus suppresses PCD without triggering plant defense responses in order to permit the biotrophic, pre-symptomatic colonization by the fungus. Therefore, the fungus may require a subtle regulation of PCD during a specific time period to help *Δbot2Δboa6#6* break through the critical point and initiate the lesion expansion phase. When overexpressing a gene with a strong promoter (e.g., the oliC promoter used in this study), the gene would be expressed at high level from the germination of conidia onwards. This would lead to CDIP accumulation during the “biotrophic stage” of the fungus, which would prematurely induce host PCD and restrict the invasion of *B. cinerea*. In order to test this hypothesis, we could overexpress gene(s) encoding PCD-inducing molecules in the WT B05.10 background and check whether these transformants trigger host resistance and result in inhibition of the invasion of the fungus.

This study illustrated that the outcome of fungal virulence assays can be strongly influenced by a (seemingly trivial) adjustment in inoculation medium. The phenomenon may be one of the causes of differences in results between different laboratories studying the same pathosystem and clearly deserves more attention in future studies.

## Materials and Methods

### Plant and fungal materials and their growth conditions

*B. cinerea* strains used and generated in this study (Table 2) were grown and conidia harvested as described (31). *S. lycopersicum* cv. Moneymaker plants were grown as described (31).

### Inoculation assays and *in vitro* cultures of *B. cinerea* for RNA sequencing

The media used were Gamborg B5 + vitamins (Duchefa, Haarlem, NL) and yeast extract (Oxoid, UK). *B. cinerea* conidia were diluted to a density of 1000/μl in Gamborg B5 media (Gamborg B5, 25 mM glucose, 10 mM potassium phosphate, pH 6.0) either without yeast extract or with 0.1 % (w/v) yeast extract. The suspension was pre-incubated for one hour before being inoculated on plants or *in vitro* cultures. For disease assays, each leaf half of *S. lycopersicum* detached leaves were inoculated with 3-4 droplets, each containing 2 μl of inoculum. Inoculated leaves were incubated in a plastic tray with a transparent lid in the lab and disease symptoms were scored at 3 dpi. The number of expanding lesions (diameter > 2mm) and non-expanding lesions (diameter ≤ 2mm) were counted for calculating disease incidence (=number of expanding lesions/total inoculations). Diameters of expanding lesions were measured by a digital caliper and leaves were photographed. Statistical analyses were performed and bar charts generated using GraphPad Prism. Details for statistics and charts are in the legends.

To sample tomato leaves for RNA-seq, the adaxial surface of leaves was inoculated in circular areas, each including five droplets containing 2 μl of inoculum. Four leaflets of one compound leaf were inoculated, and one leaflet was sampled at each time point (t = 0, 12, 16 and 24 hours post inoculation (hpi)). Tomato leaves were mock-inoculated with the -yeast extract medium and collected at 0, 12, 16 and 24 hpi. Eight ml of *B. cinerea* conidia suspensions in two media (-/+ yeast extract) were pipetted on glass petri-dishes (90 mm) and incubated in the lab. Fungal tissue was sampled at 0, 12, 16 and 24 hpi for *in vitro* samples. Three biological replicates were collected for all infected and mock-inoculated tomato leaf samples (Table S1). Two biological replicates were collected for *B. cinerea in vitro* cultures (Table S1). Samples were freeze-dried, RNA was extracted using a Maxwell® 16 LEV Plant RNA Kit (Promega).

### RNA sequencing and identification of differentially expressed genes (DEGs)

Strand-specific libraries of RNA samples were constructed, followed by paired-end sequencing on a DNBseq platform (BGI Tech Solutions, Hongkong), with a read depth of 20-50 Million (Table S1). Mapping and quantifying gene transcripts from RNA-seq reads were performed using Kallisto (v0.44.0) (32), with a 100 bootstrap value. Principal component analysis (PCA) was conducted as transcriptome samples clustered by the ggplot2 package in R. Sleuth (v0.30.0) was used for differential expression analysis (33). DEG analysis was done with default settings, removing genes that have <5 estimated read counts in >47% of all the samples. Genes were considered differentially expressed if they displayed between two samples a log2 fold change ≥ 2 or ≤ −2 with an adjusted p-value ≤ 0.05 (Benchamini–Hochberg method). UpSet plots were generated using R package UpSetR (34).

### Network construction and co-expression analysis

80 samples were used for constructing co-expression analysis using R package WGCNA (v1.69) (35). The normalized gene expression matrix was imported into WGCNA to construct co-expression modules. The gene expression matrix was searched for a suitable soft threshold to build gene networks using a scale-free topology model (36). The scale-free network obtained by power processing at β = 13, resulted in an adequate fit with r^2^ = 0.85 with average connectivity approaching 0. Therefore, β = 13 was used to construct a scale-free network. The adjacency matrix was transformed into a topological overlap matrix (TOM) to evaluate the correlation between expression profiles of genes (36). The dissimilar topological matrix (dissTOM= 1-TOM) was used to carry out matrix clustering and module partitioning by the dynamic shearing algorithm. The minimum number of elements in a module was 30 (minModule Size=30), and the threshold for merging of a similar module 0.25 (CutHeight = 0.25) (Fig. S5). Module eigengenes (MEs) were used to calculate correlation coefficients to traits to identify the biologically significant modules. The Pearson correlation coefficient of the corPvalueStudent () function was used to calculate correlations between the infection and yeast extract matrix and the module feature gene matrix to obtain p-values. Both a positive and negative correlation can suggest involvement in either response and we chose thresholds of [r] > 0.30 and p < 0.01 as being significant.

### Gene Ontology analysis of module genes

Genes in a module were mapped to terms in the Gene Ontology database (http://www.geneontology.org/), and were enriched as compared to the genome background (37). The GO terms were subsequently filtered for the ones belonging to the domains of biological processes and molecular functions. Numbers were calculated for every term, and enriched GO terms in modules were defined by hypergeometric distribution algorithm. FDR was set to a threshold ≤ 0.05. GO enrichment was visualized using ggplot2 (R package) (38).

### *B. cinerea* transformation by CRISPR-Cas9 mediated approach

*B. cinerea* mutant strains were generated by CRISPR-Cas9 mediated transformation with minor modification from (39). Primers for synthesis of sgRNAs and amplification of donor templates are in Table S2. *Bcssp2, Bcpg1* and *Bcnep2* were amplified from B05.10 genomic DNA by PCR using Phusion Hot Start II DNA Polymerase (Thermo Scientific™) and cloned into pNDH-OGG or pNAN-OGG (40) by replacing the GFP cassette using the ClonExpress MultiS One Step Cloning Kit (Vazyme). Donor templates for transformation were amplified from pNDH-OGG harboring *Bcssp2*, or from pNAN-OGG vector harboring either *Bcpg1* or *Bcnep2*. *Bcssp2* was overexpressed by replacing *BcniaD* in the *Δbot2Δboa6#6* recipient strain via homologous recombination upon cleavage of Cas9 at *BcniaD* to obtain *ΔΔ-OEssp2* transformants, while *Bcpg1* or *Bcnep2* was overexpressed by replacing *BcniiA* to obtain *ΔΔ-OEpg1* or *ΔΔ-OEnep2* mutants. Methods for molecular characterization of transformants are described in (31) using primers in Table S2.

## Acknowledgements

The research of Si Qin, Yaohua You and Jie Chen was funded by the China Scholarship Council We are grateful to prof. Matthias Hahn (University Kaiserslautern, Germany) for providing the *B. cinerea* mutant strains *Δbot2#1, Δbot2Δboa6#6*, 12xbb, 12xpg and 18x, to dr. Muriel Viaud (INRAE-BIOGER, France) for providing the *Δboa6#1* mutant and to dr. Julia Schumacher (BAM, Berlin, Germany) for providing the pNDH-OGG and pNAN-OGG plasmids.

## Data availability

Sequence data have been deposited in the NCBI Sequence Read Archive (SRA). The accession numbers are pending.

## Supplementary material

**Fig. S1.**
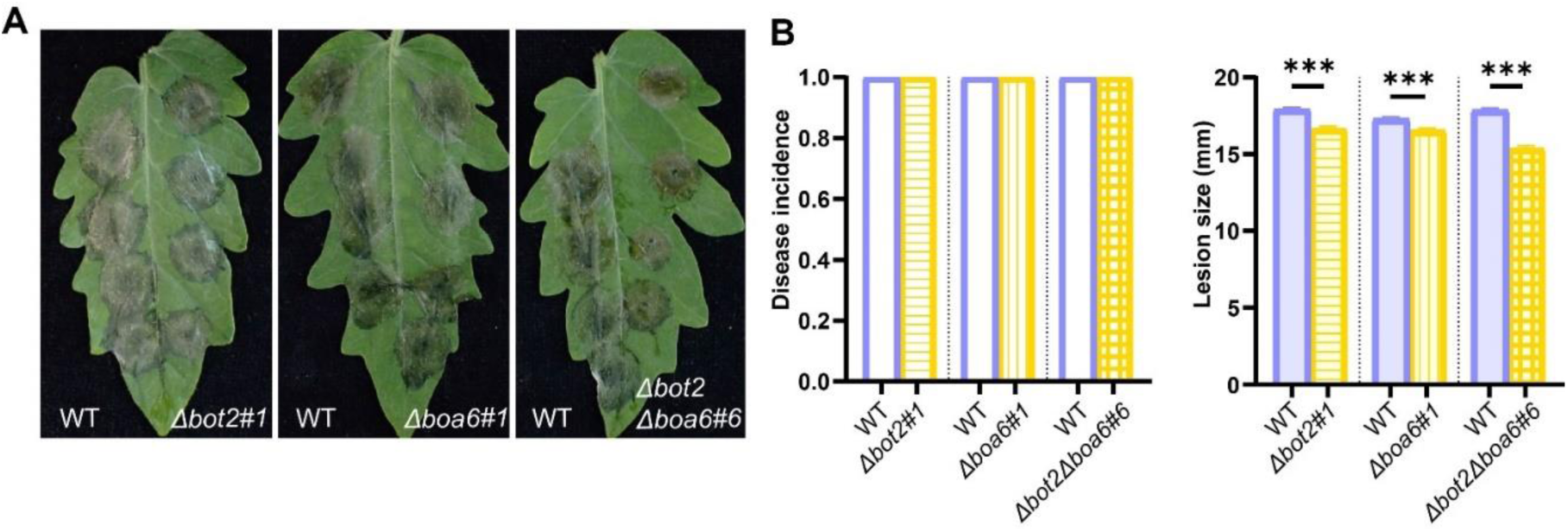
Virulence of *Δbot2#1, Δboa6#1* and *Δbot2Δboa6#6* compared with WT B05.10 on tomato leaves, examined by infection assays performed with PDB as inoculation medium. **(A)** The symptoms on tomato leaves infected by each mutant strain (on the right of the central vein) compared with WT B05.10 (on the left of the central vein), photographed at 3 dpi. **(B)** Bar charts of disease incidences (left) and lesion sizes measured by a digital caliper (right) at 3 dpi, presented as means with standard errors from 96 datapoints from three independent experiments. Asterisks indicate the significant differences (***p<0.001) by t-test.

**Fig. S2.**
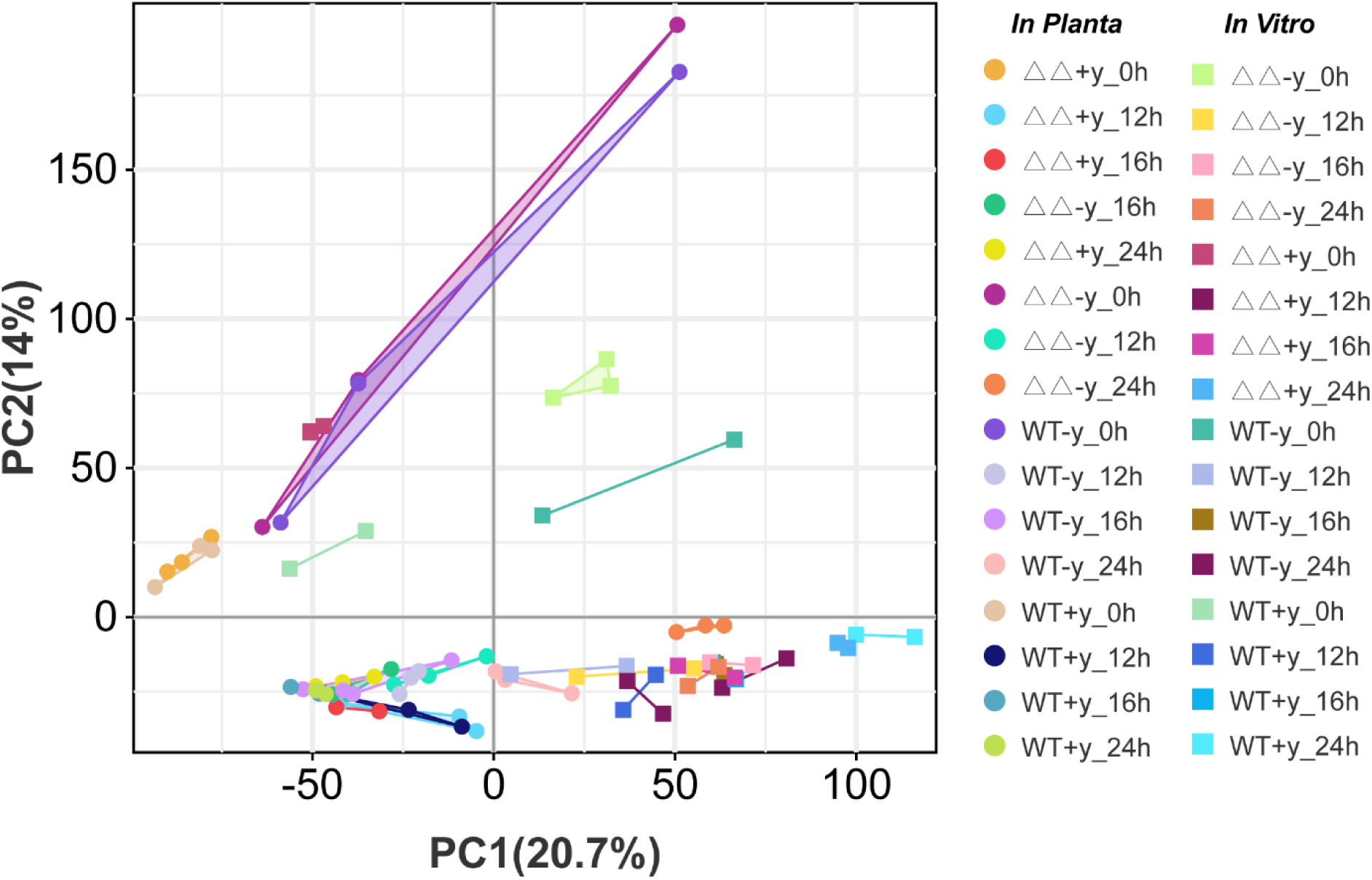
PCA plot of the RNA-seq dataset shows the differences (distances) between all samples generated in this study, according to gene expression levels. Each round dot indicates an *in planta* sample, and each squared dot indicates an *in vitro* sample. Different colors represent different fungal strains (WT B05.10 and *Δbot2Δboa6#6* (WT and ΔΔ)), different media (- and +y), and different timepoints (0, 12, 16 or 24 hpi).

**Fig. S3.**
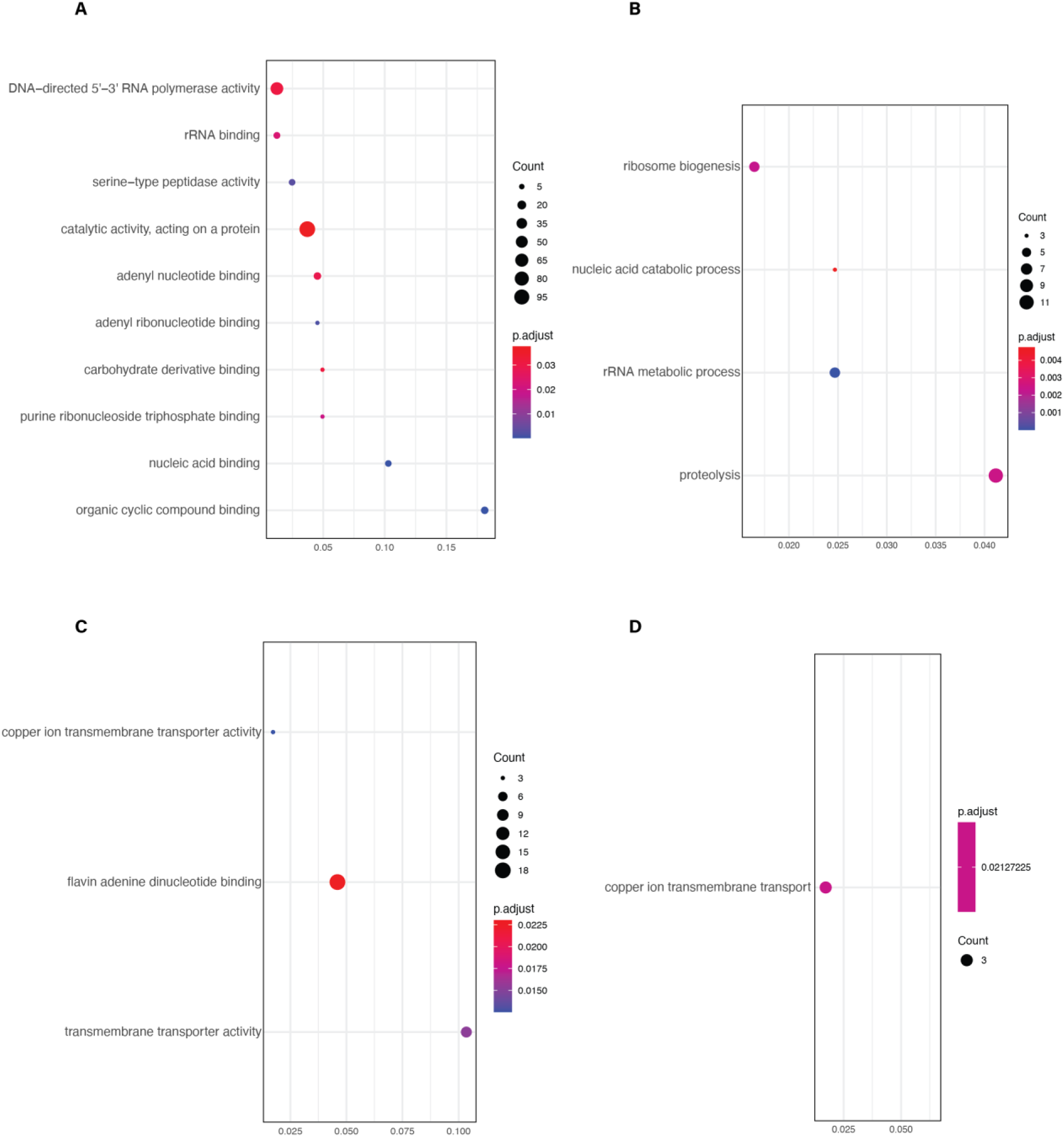
GO enrichment analysis GO enrichment analysis for genes that are up-regulated between +yeast and -yeast (A and B) and down-regulated (C and D) both in WT and mutant at 24 hpi. (A, C) Molecular function. (B, D) Biological process.

**Fig. S4.**
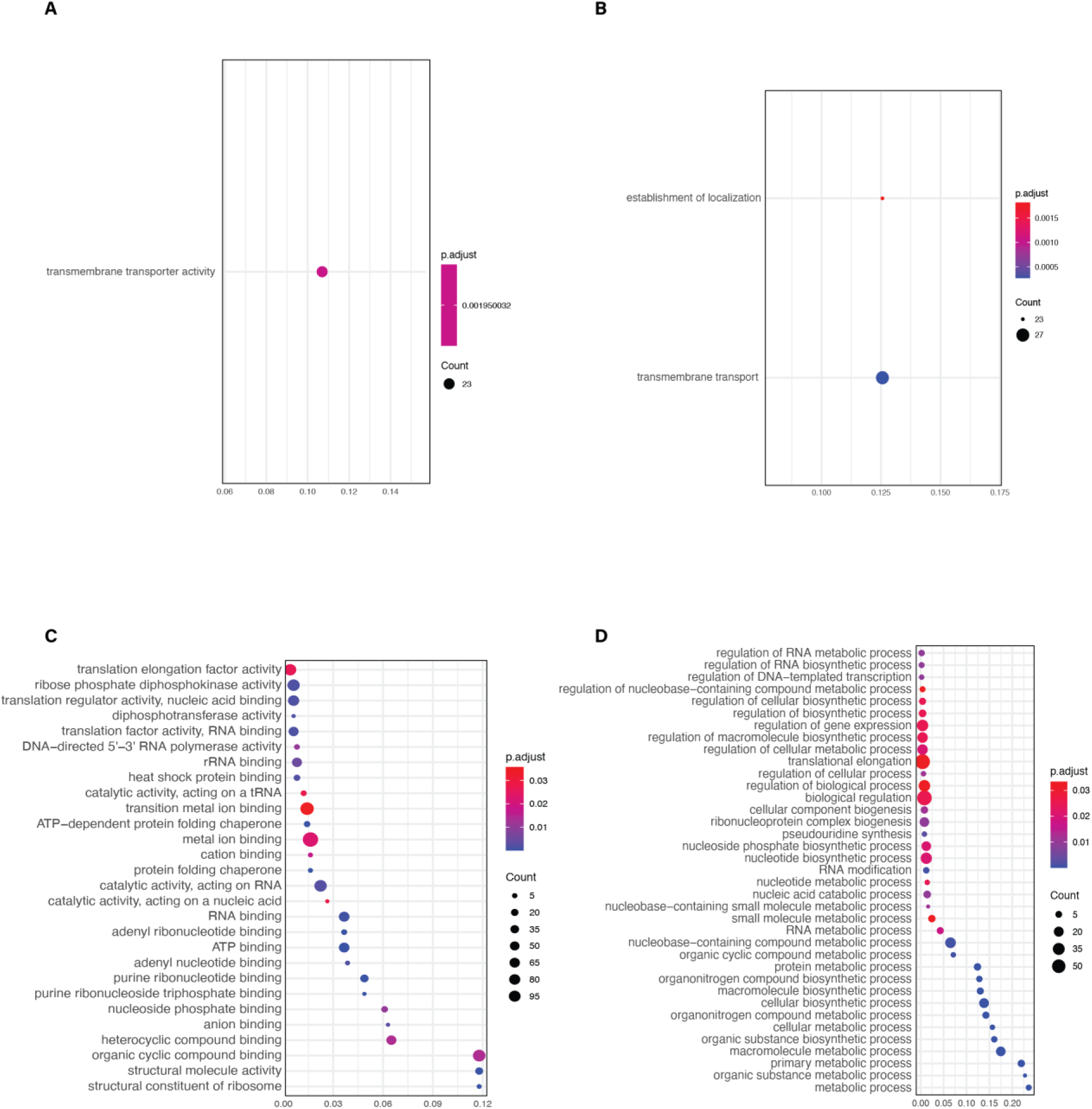
GO enrichment analysis GO enrichment analysis for DEGs (+yeast vs -yeast) that are unique in WT (A, B) and mutant (C, D) at 24 hpi. WT only contains enriched GO terms in down-regulated DEGs (A, B), whereas mutant contains only GO terms in up-regulated genes (C, D). (A, C) Molecular function. (B, D) Biological process.

**Fig. S5.**
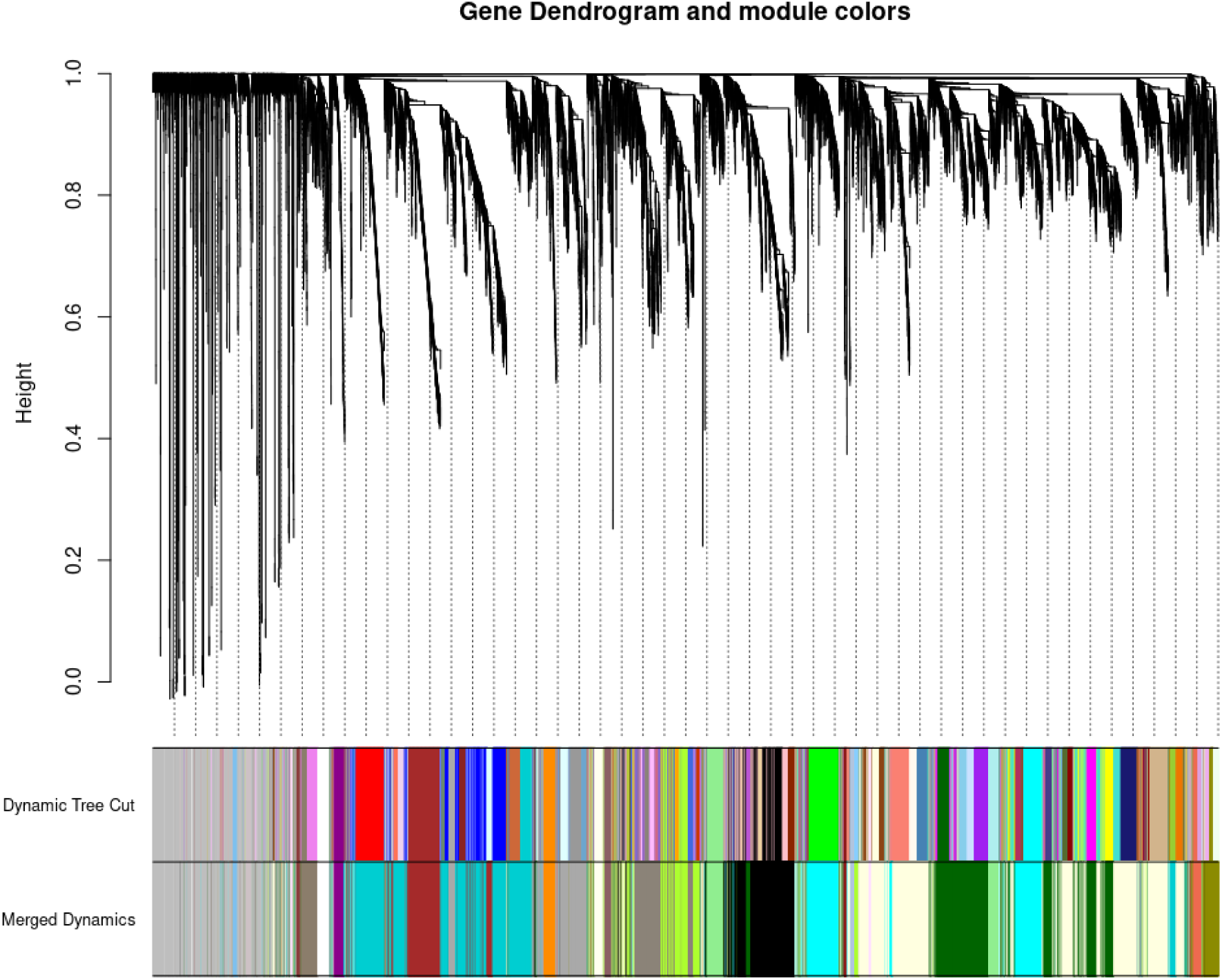
Dendrogram of gene co-expression network. In the top part of the figure, a dendrogram of the filtered set of genes is shown, constructed using the topological overlap between genes. Below the dendrogram, the top colored bar indicates the co-expression modules produced by the dynamic tree cutting algorithm, and the bottom colored bar indicates the merged co-expression modules.

**Fig. S6.**
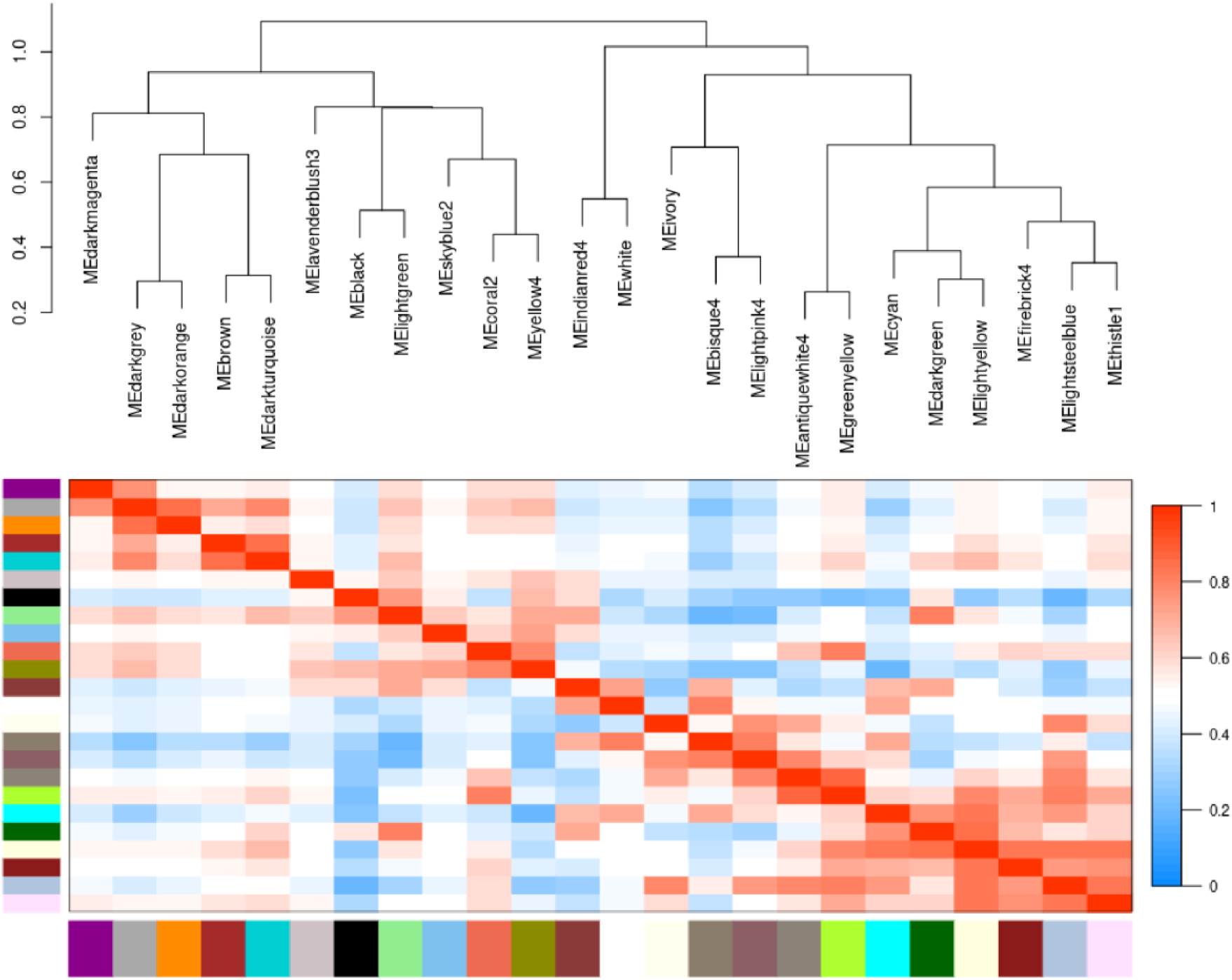
Module significance correlation based on the Pearson correlation.

**Fig. S7.**
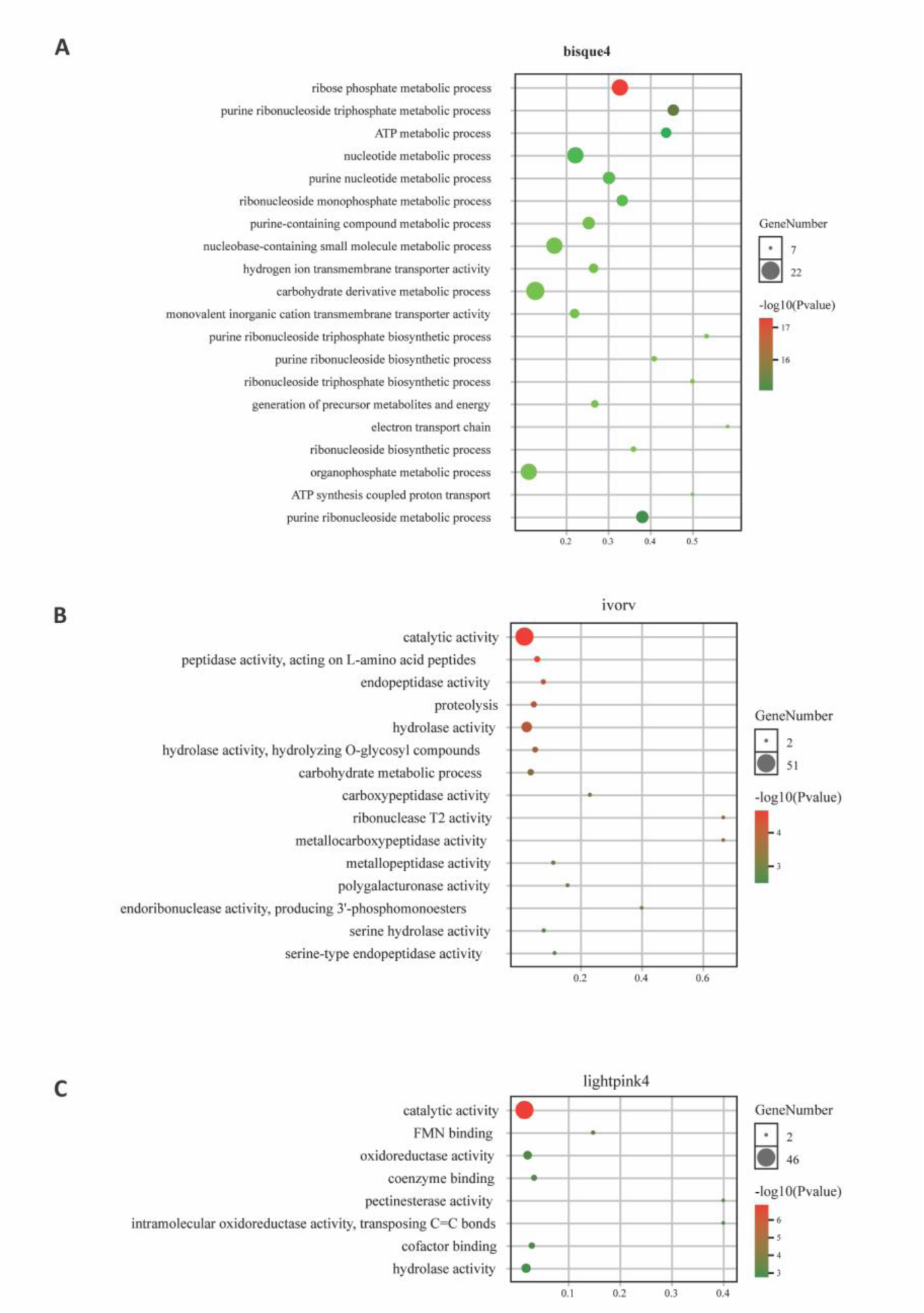
GO enrichment analysis of three gene modules GO enrichment analysis of three gene modules that are positively corrected to compatible interaction between *B. cinerea* and tomato.

**Fig. S8.**
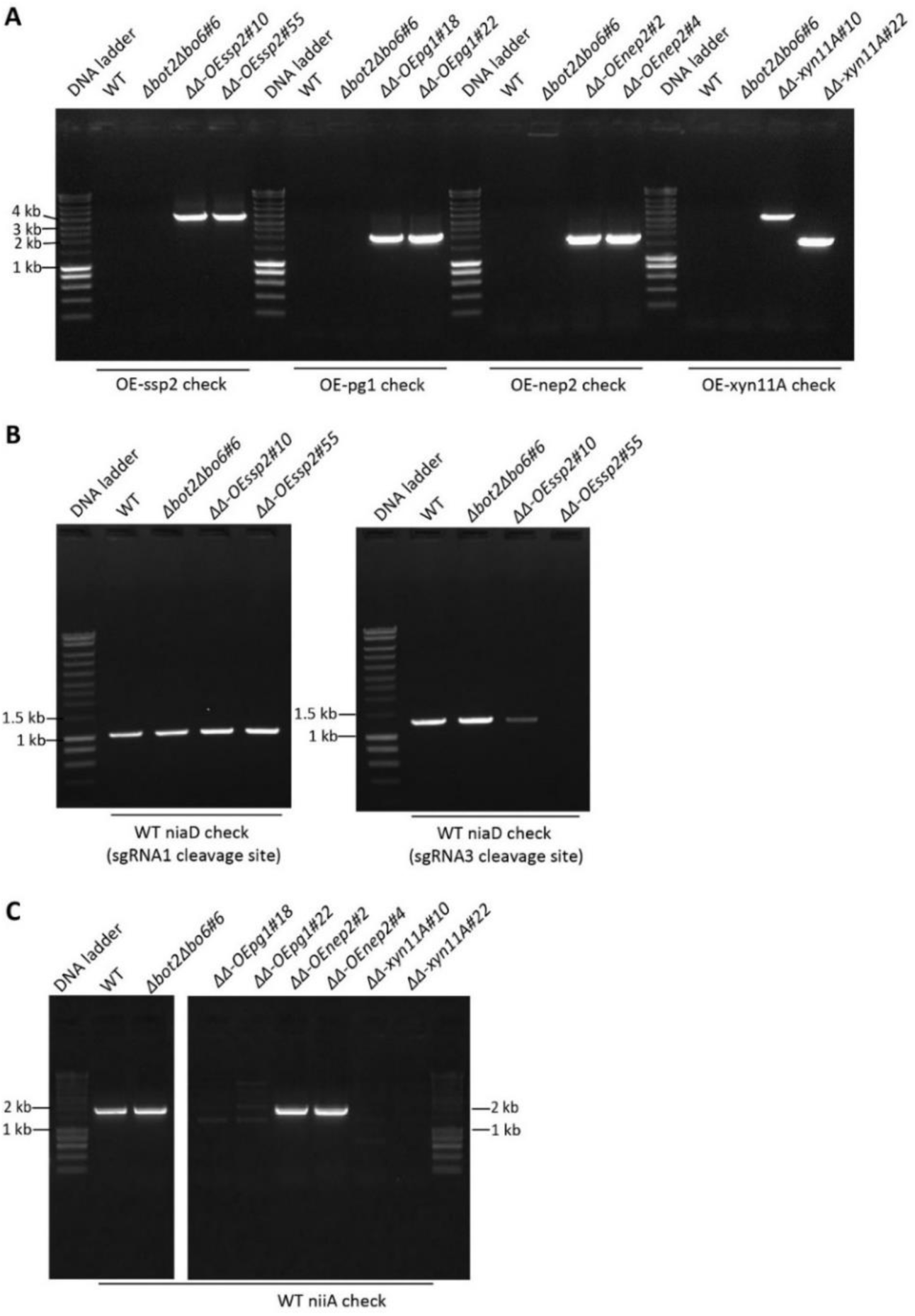
Molecular characterization of overexpression mutants Molecular characterization of overexpression mutants *ΔΔ-OEssp2#10, ΔΔ-OEssp2#55*, *ΔΔ-OEpg1#18*, *ΔΔ-OEpg1#22, ΔΔ-OEnep2#2, ΔΔ-OEnep2#4, ΔΔ-OExyn11A#10* and *ΔΔ-OExyn11A#22*. WT B05.10 and *Δbot2Δboa6#6* double mutant were used as controls. **(A)** PCR results confirmed that integration of the overexpression cassettes for *ssp2*, *pg1*, *nep2* and *xyn11A* were successful. Two independent transformants for each construct were obtained. The donor-templates for the OE*ssp2* and *OExyn11A* transformants were inserted ectopically inside of the *niaD* gene instead of replacing the whole *niaD* coding sequence. A similar strategy was chosen for the. The other transformants were obtained via replacing the entire *niiA* gene by the corresponding overexpression*-*donor-template. **(B)** The insertion of OE*ssp2-*donor-template by NHEJ was confirmed to be located at the sgRNA3 cleavage site for *ΔΔ-OEssp2#55* which was a homokaryotic transformant. *ΔΔ-OEssp2#10* was either a heterokaryon or the ectopic insertion location cannot be confirmed by these PCR. (C) *ΔΔ-OEpg1#18*, *ΔΔ-OEpg1#22, ΔΔ-OExyn11A#10* and *ΔΔ-OExyn11A#22* were confirmed to be homozygous transformants, while *ΔΔ-OEnep2#2* and *ΔΔ-OEnep2#4* were heterokaryons. Names of the *B. cinerea* strains are indicated above each DNA gel picture, and the aims for the primer combinations are indicated below the gel.

**Table S1.** Overview of all samples in this study that were used for sequencing.

| Sample_name | Strain | Inoculation<br>medium | *In planta* (# of<br>replications) |  |  |  | *In vitro* (# of<br>replications) |  |  |  |
| --- | --- | --- | --- | --- | --- | --- | --- | --- | --- | --- |
|  |  |  | 0 | 12 | 16 | 24 | 0 | 12 | 16 | 24 |
|  |  |  | hpi | hpi | hpi | hpi | hpi | hpi | hpi | hpi |
| WT-y | B05.10 | - y<sup>1</sup> | 3 | 3 | 3 | 3 | 2 | 2 | 2 | 2 |
| WT+y | B05.10 | + y<sup>2</sup> | 3 | 3 | 3 | 3 | 2 | 2 | 2 | 2 |
| $\Delta\Delta$-y | $\Delta bot2\Delta boa6\#6$ | - y | 3 | 3 | 3 | 3 | 3 | 2 | 2 | 2 |
| $\Delta\Delta$+y | $\Delta bot2\Delta boa6\#6$ | + y | 3 | 3 | 3 | 3 | 3 | 2 | 2 | 2 |

<sup>1</sup> Gamborg B5, 25 mM glucose, 10 mM potassium phosphate, pH 6.0; <sup>2</sup>Gamborg B5, 25 mM glucose, 10 mM potassium phosphate, 0.1 % yeast extract, pH 6.0

**Table S2.**
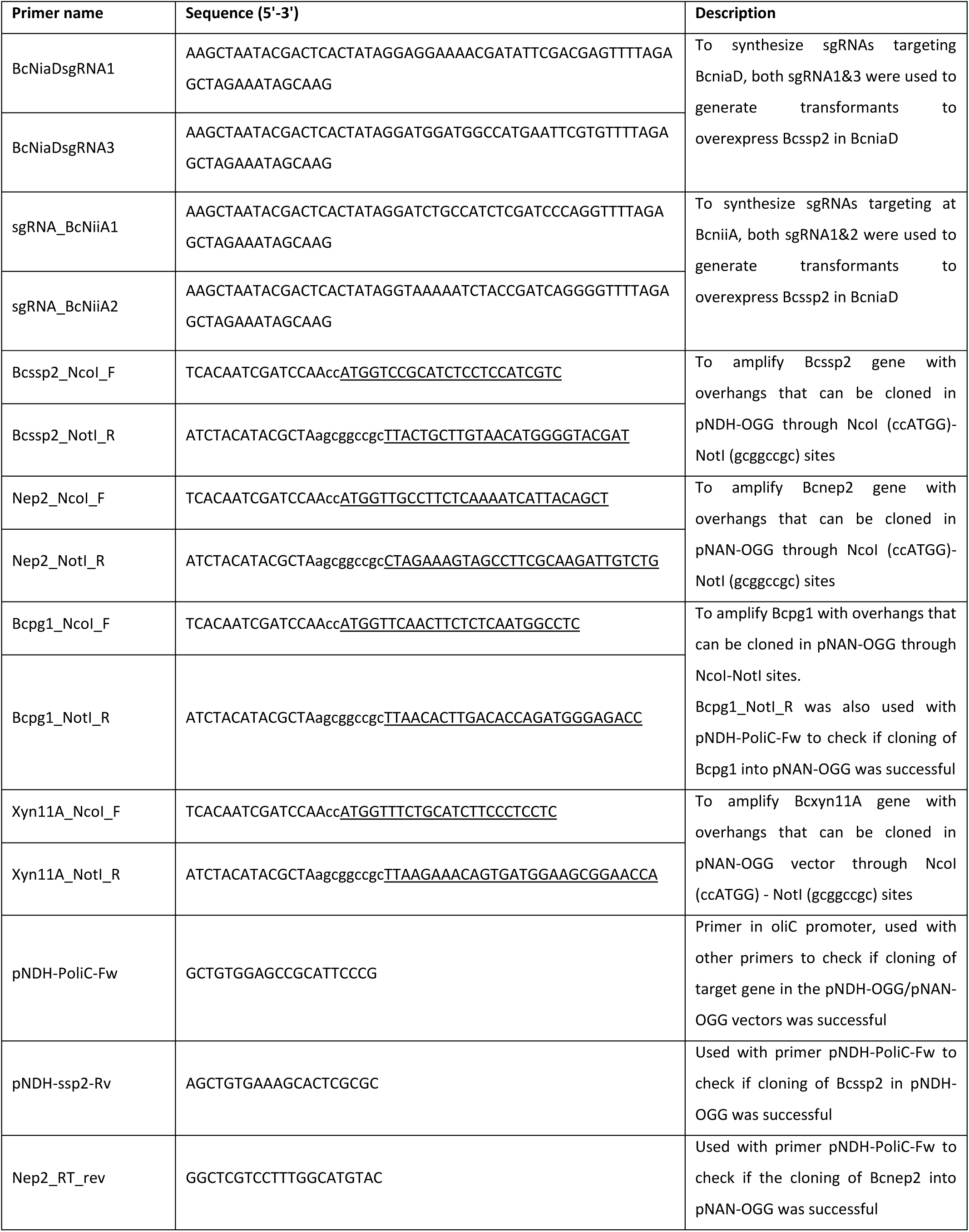
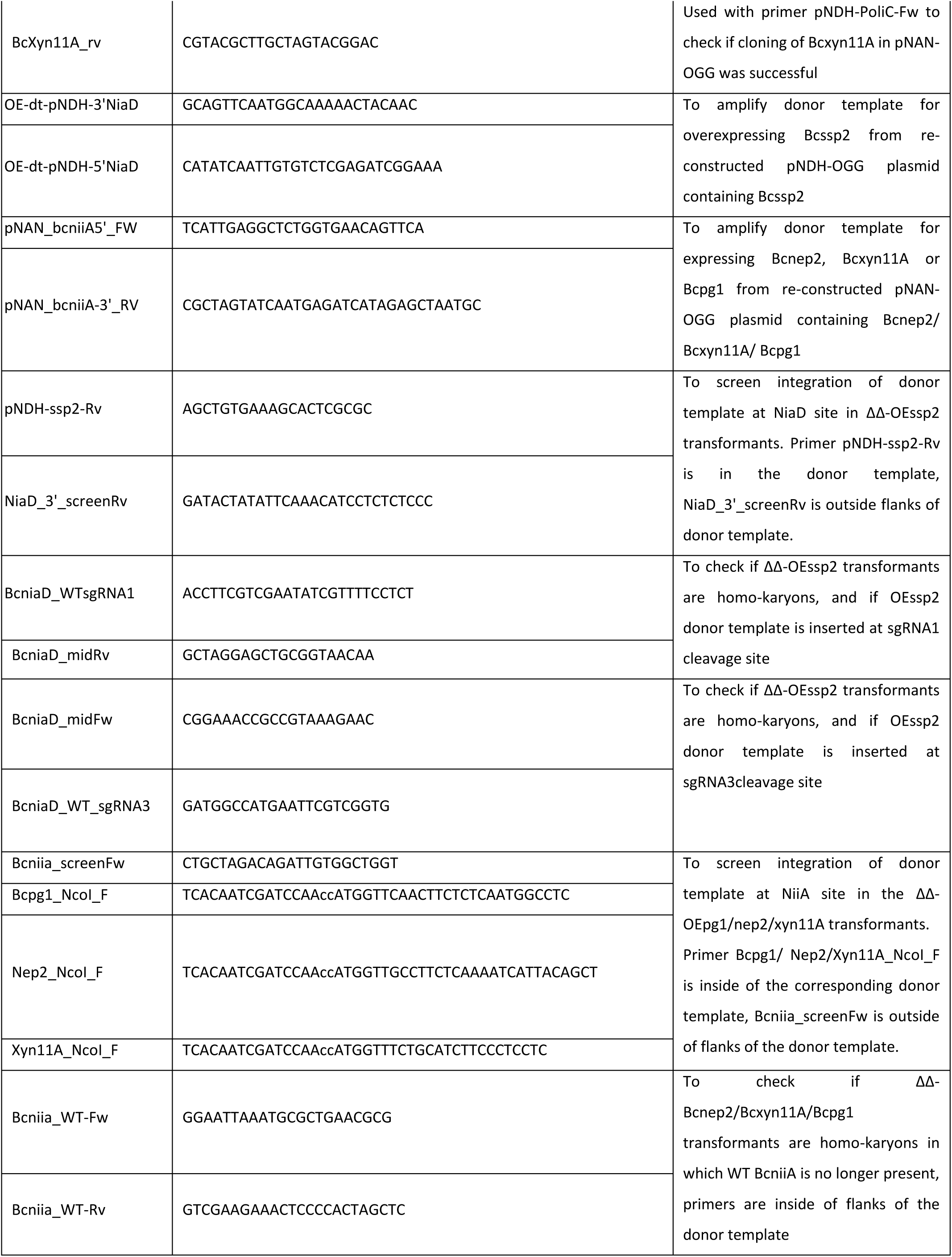
Primers used in this study.

## Legend supplementary data

**Supplementary Data S1.** Genes encoding secreted proteins in three modules are positively correlated to the compatible interaction. Sheet 1 (named “Summary”) contains an overview of the total number of genes encoding secreted proteins in each module resulting from the GGCNA analysis. Sheets 2-4 contains gene lists for module “lightpink4”, “bisque4”, and “ivory”, respectively.

